# Coevolution of the olfactory organ and its receptor repertoire in ray-finned fishes

**DOI:** 10.1101/2021.12.13.472396

**Authors:** Maxime Policarpo, Katherine E Bemis, Patrick Laurenti, Laurent Legendre, Jean-Christophe Sandoz, Sylvie Rétaux, Didier Casane

**Affiliations:** Université Paris-Saclay, CNRS, IRD, UMR Évolution, Génomes, Comportement et Écologie, 91198, Gif-sur-Yvette, France; NOAA National Systematics Laboratory, National Museum of Natural History, Smithsonian Institution, Washington, D.C. 20560, U.S.A; Université de Paris, Laboratoire Interdisciplinaire des Energies de Demain, Paris, France; Université Paris-Saclay, CNRS, Institut des Neurosciences Paris-Saclay, 91400 Saclay, France; Université de Paris, UFR Sciences du Vivant, F-75013 Paris, France

**Keywords:** olfactory epithelium, olfactory lamellae, olfactory receptor genes, Actinopterygii, gene family dynamics

## Abstract

Ray-finned fishes (Actinopterygii) perceive their environment through a range of sensory modalities, including olfaction ^1,2^. Anatomical diversity of the olfactory organ suggests that olfaction is differentially important among species ^1,3,4^. To explore this topic, we studied the evolutionary dynamics of the four main gene families (OR, TAAR, ORA/VR1 and OlfC/VR2) ^5^ coding for olfactory receptors in 185 species of ray-finned fishes. The large variation in the number of functional genes, between 28 in the Ocean Sunfish *Mola mola* and 1317 in the Reedfish *Erpetoichthys calabaricus*, is the result of parallel expansions and contractions of the four main gene families. Several ancient and independent simplifications of the olfactory organ are associated with massive gene losses. In contrast, polypteriforms, which have a unique and complex olfactory organ, have almost twice as many olfactory receptor genes as any other ray-finned fish. These observations suggest a functional link between morphology of the olfactory organ and richness of the olfactory receptor repertoire. Further, our results demonstrate that the genomic underpinning of olfaction in ray-finned fishes is heterogeneous and presents a dynamic pattern of evolutionary expansions, simplifications and reacquisitions.

## Introduction

With more than 34,000 valid species, Actinopterygii (ray-finned fishes) is the largest group of aquatic vertebrates ^6^. Most species of ray-finned fishes belong to Teleostei (teleosts), but a few extant species belong to relictual clades: Polypteriformes, Acipenseriformes, Lepisosteiformes and Amiiformes (**Fig. 1**). With a last common ancestor that lived 368-379 Million years ago (Ma) ^7,8^, the remarkable taxonomic diversity of actinopterygians comes with striking anatomical, physiological, behavioral and ecological adaptations ^9^.

**Fig. 1.**
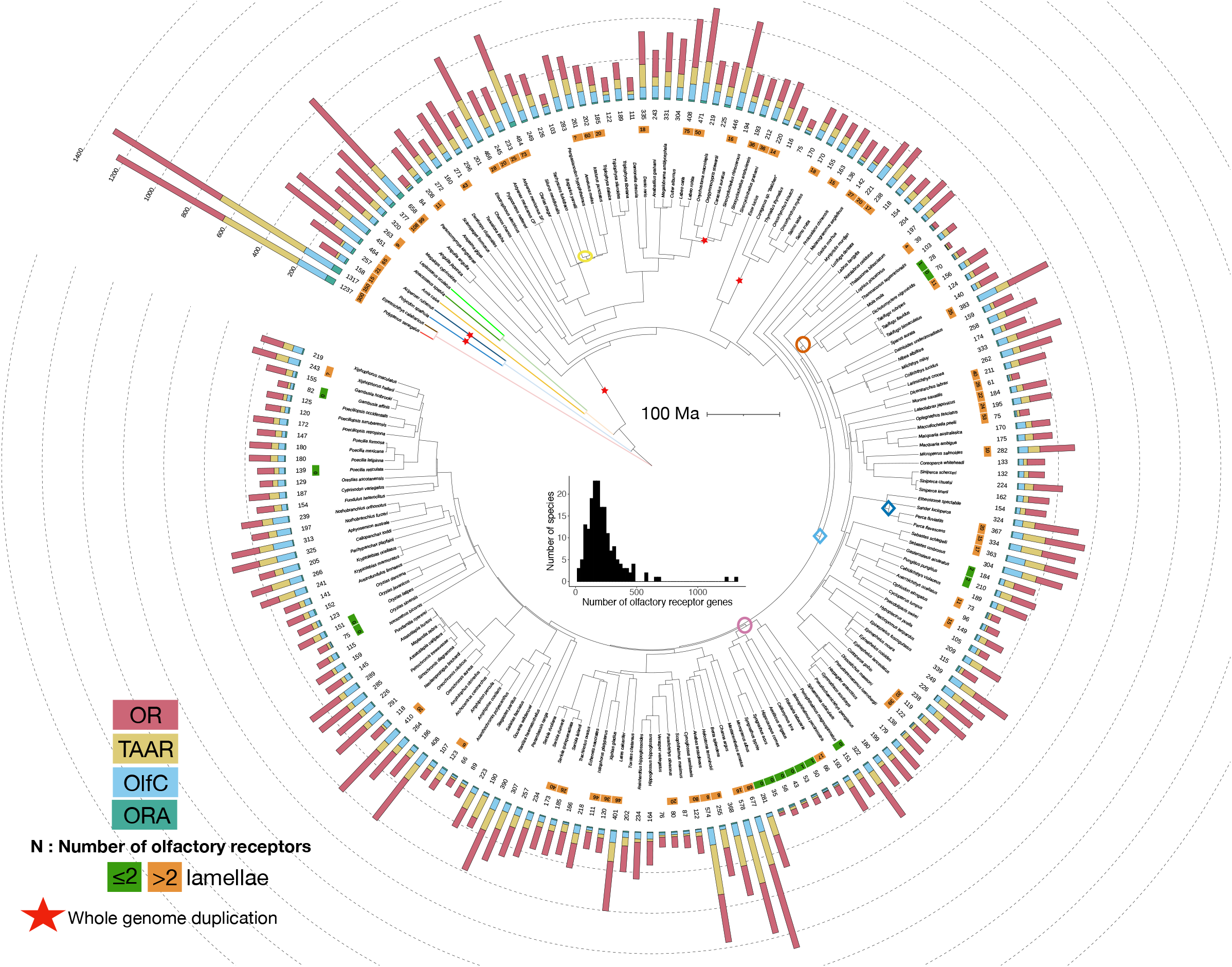
Diversity of olfactory receptor gene repertoire in ray-finned fishes (Actinopterygii). Time-calibrated phylogeny from https://fishtreeoflife.org/. For each species, a barplot represents the number of OR, TAAR, OlfC and ORA genes. Where available, number of olfactory lamellae is indicated. Branches associated with two highest birth rates and two highest death rates are indicated by diamond and oval symbols, respectively. Branch color code: Polypteriformes (red, *Polypterus senegalus*; brown *Erpetoichthys calabaricus*); Acipenseriformes (light blue *Polyodon spathula*; dark blue *Acipenser ruthenus*); Amiiformes (yellow *Amia calva*); Lepisosteiformes (dark green *Atractosteus spatula*; light green *Lepisosteus oculatus*); teleost (black). The phylogeny was visualized using iTOL. Distribution of the total number of olfactory receptor genes per species is shown in center of figure.

Actinopterygians thrive in aquatic habitats from the tropics to the polar regions, in small temporary ponds to large oceans.

Ray-finned fishes have several sensory systems to process physical and chemical cues. Among them, the olfactory system serves in feeding, reproduction, predator avoidance and migration ^10^. In a seminal work, Burne (1909) ^3^ described the great anatomical diversity in the olfactory organs of ray-finned fishes. Since then, it has been assumed that fishes with a multilamellar olfactory epithelium have a better sense of smell than those with a flat olfactory epithelium, respectively classified as macrosmatic and microsmatic ^11^.

In most ray-finned fishes, the olfactory epithelium forms a rosette in which lamellae attach to a central raphe (e.g., *Danio rerio* in **Fig. 2A**). Chondrichthyans (sharks, rays, chimaeras) also have olfactory rosettes ^12^, thus it is likely that olfactory rosettes were present in the common ancestor of jawed vertebrates and conserved in the common ancestor of ray-finned fishes. However, the olfactory rosette has been simplified several times during the evolution of ray-finned fishes, leading in the most extreme cases to a small, flat olfactory epithelium with no lamellae (e.g., *Syngnathus typhle* in **Fig. 2A**) ^4,13^. In contrast, other groups have multilamellar organizations of the olfactory epithelium. The most extreme example of a multilamellar olfactory epithelium occurs in the Polypteriformes, which have a large and complex structure: a nasal capsule is divided into six sectors, five in a main sac and one in a diverticulum, each with a rosette-like organization with a septum and lamellae attached to both sides (e.g., *Polypterus senegalus* and *Erpetoichthys calabaricus* in **Fig. 2A**) ^14,15^.

**Fig. 2.**
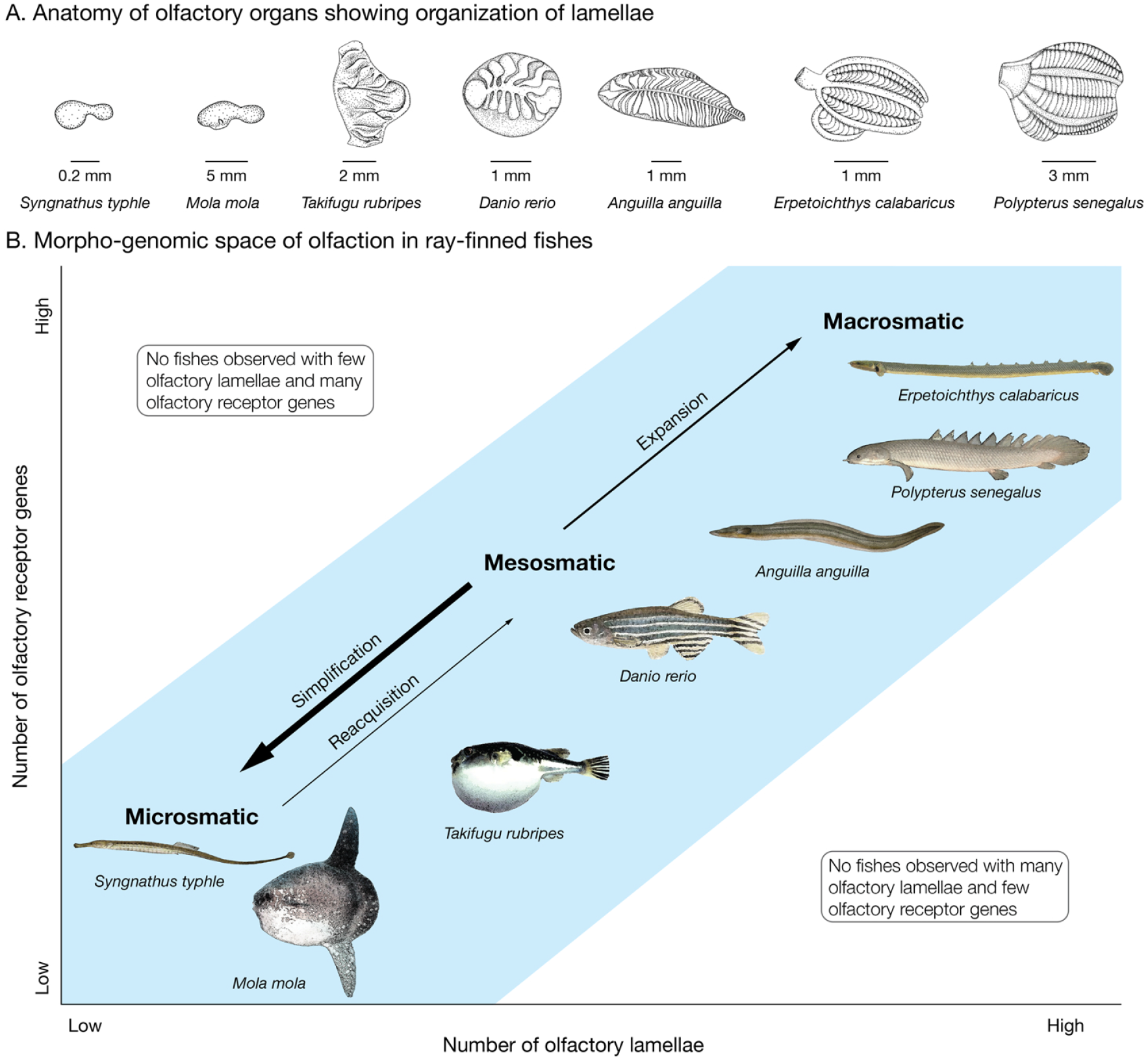
Morpho-genomic space of olfaction in ray-finned fishes. (A) Diversity of olfactory organ morphology. *Syngnathus typhle*, 283 mm TL, *Mola mola*, 1290 cm TL, *Takifugu rubripes* 290 mm TL, *Danio rerio*, 30 mm TL, *Anguilla anguilla*, 450 mm TL, *Erpetoichthys calabaricus*, 268 mm TL, *Polypterus senegalus*, 112 mm TL. Anterior to left. (B) Correlation between number of olfactory lamellae and number of olfactory receptor genes; all fishes examined, ranging from microsmatic to macrosomatic, occurred in the blue region of the graph. Most evolutionary transitions in the olfactory organ, indicated by arrows, were simplifications (e.g., *S. typhle, M. mola*), but expansions (e.g., *A. anguilla, E. calabaricus*, and *P. senegalus*) and reacquisition (e.g., *T. rubripes*) also occurred.

The diversity of odorants that can be detected depends on the size of the olfactory receptor gene repertoire ^16^. In vertebrates, olfactory receptor genes belong to four main gene families with independent origins: odorant receptors (OR), trace amine-associated receptors (TAAR), and vomeronasal receptors 1 and 2 (named VR1 and VR2 in tetrapods). Actinopterygian fishes do not have a vomeronasal organ, and thus VR1 and VR2 gene families are referred to as ORA and OlfC, respectively. Only a few olfactory receptor genes have been identified that do not belong to these four gene families ^17^.

Analyses of genomes of 13 teleosts and one non-teleost suggested that ORA is a small and stable gene family, with eight genes in the last common ancestor of ray-finned fishes and six genes in most teleosts ^18^. More genes have been identified in OR, TAAR and OlfC gene families ^19–21^, but only the evolutionary dynamics of OR genes have been analyzed broadly in teleosts ^13^. Policarpo et al. (2021)^13^ found a ~30-fold variation in the number of OR genes (15 in Ocean Sunfish *Mola mola* and Broad-nose Pipefish *Syngnathus typhle* to 429 in Zig-zag Eel *Mastacembelus armatus*) and reported that the number of olfactory lamellae correlates with the richness of the OR gene repertoire.

A recent burst of high quality ray-finned fish genomes, in particular for non-teleost actinopterygians such as polypteriforms, prompted us to analyze the evolution the four olfactory gene families and the anatomy of the olfactory organ across the phylogeny of ray-finned fishes.

## Results/Discussion

### Coevolutionary dynamics of olfactory gene families

We characterized the olfactory receptor gene repertoire, including OR, TAAR, OlfC and ORA genes, for 185 species of ray-finned fishes selected on the basis of high genome completeness (**Fig. 1, Extended Data Fig. 1 and Supplementary data 1)**.

The mean size of the total olfactory gene repertoire for actinopterygians was 224 genes. The largest (1317 genes) was found in the polypteriform *Erpetoichthys calabaricus* (Reedfish) and the smallest (28 genes) in the tetraodontiform *Mola mola* (Ocean Sunfish) (**Fig. 1**).

ORA is a small and stable family typically comprising six genes (ORA1 to ORA6) in teleosts ^18^. Nevertheless, we found up to three ORA genes have been lost in several lineages, and, surprisingly, this gene family is much larger in some lineages, particularly polypteriforms, which have nearly 50 functional ORA genes (**Fig. 1** and **Extended Data Fig. 1A**). Two genes, ORA7 and ORA8, were present in the last common ancestor of ray-finned fishes; ORA7 was lost in the common ancestor of teleosts, while ORA8 was lost in clupeocephalans (**Extended Data Fig. 2**).

The evolution of the other three gene families (OR, TAAR, OlfC) has been more dynamic. For example, we identified an average of 126 functional OR genes in ray-finned fishes, but the variance is large, with 623 and 606 OR genes in the polypteriforms *Erpetoichthys calabaricus* and *Polypterus senegalus*, respectively, and only 15 OR genes in the Ocean Sunfish *Mola mola* and Broad-nose Pipefish *Syngnathus typhle* (**Fig. 1** and **Extended Data Fig. 1B**). The OR family is split into seven monophyletic subfamilies, α, β, γ, δ, ε, and η ^22^. In tetrapods, α and γ families expanded and other subfamilies are relictual or absent. In contrast, in teleosts the α family is absent and only one copy of a γ family gene occurs in Zebrafish *Danio rerio* ^22^. Our analysis shows that α family genes occur in all non-teleost actinopterygians, but that the α family was lost in the common ancestor of teleosts (**Extended Data Fig. 1B**). The γ family is well represented in non-teleost actinopterygians whereas only a few copies are scattered in the teleost phylogeny (**Extended Data Fig. 1B**). This suggests that the γ family was present in the common ancestor of teleosts but lost in most teleost lineages.

The number of genes in the TAAR and OlfC repertoires is smaller than in the OR repertoire, with an average of 51 and 40 genes per species, respectively. For these two gene families, the variance is also large. For example, the polypteriform *Erpetoichthys calabaricus* has 486 TAAR and 161 OlfC genes. At the opposite extreme, only three TAAR genes were found in the syngnathiform *Callionymus lyra* and two OlfC genes in tetraodontiform *Mola mola* (**Fig. 1** and **Extended Data Fig. 1C,D**).

To analyze the evolutionary dynamics of the olfactory receptor gene families, we computed birth and death rates along branches of the phylogeny for OR, TAAR and OlfC families using the gene tree – species tree reconciliation method ^23^. The mean birth and death rates were similar in OR, TAAR and OlfC families, 0.0071/0.0071, 0.0101/0.0079 and 0.0059/0.0069 per gene per million years, respectively (**Extended Data Fig. 3)**. Whereas birth and death rates are similar along most branches, we observed concomitant high death rates of OR, TAAR and OlfC genes in the common ancestor of two sampled species of Siluriformes (*Bagarius yarrelli* and *Tachysurus fulvidraco*), in the common ancestor of Lophiiformes and Tetraodontiformes, and in the common ancestor of Kurtiformes and Syngnathiformes. We also observed concomitant high birth rates of OR, TAAR and OlfC genes in the common ancestor of Labriformes and Cyprinodontiformes, and in the common ancestor of *Perca* + *Sander* (**Fig. 1**, **Extended Data Fig. 3**).

Despite variation in the number of genes in a family, we did not find evidence that contraction of one gene family is compensated by expansion of others. On the contrary, there is a correlation between the number of functional genes in each family (phylogenetic generalized least squares (PGLS); R^2^ = 0.50 between OR and TAAR, R^2^ = 0.56 between OR and OlfC, R^2^ = 0.40 between TAAR and OlfC, all p-values < 2e-16, **Fig. 3**).

**Fig. 3.**
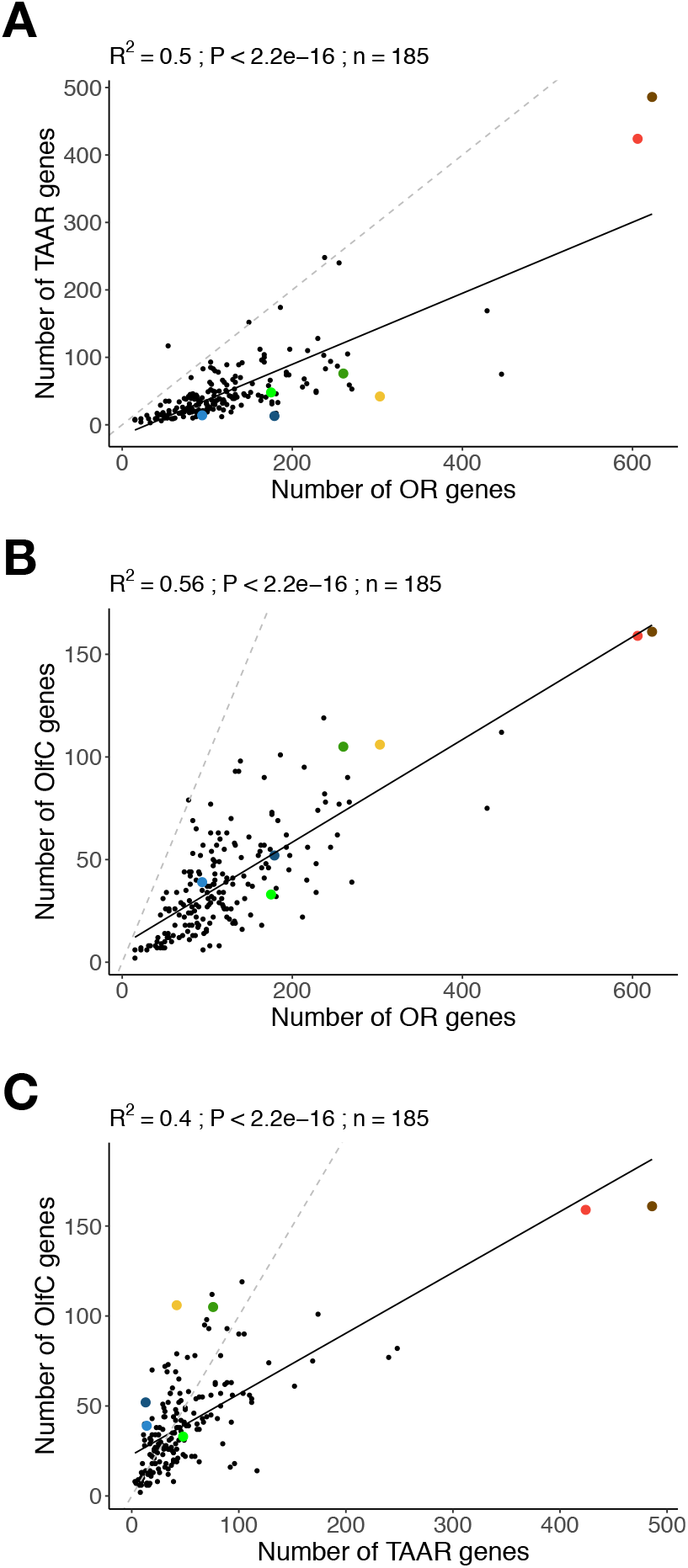
Coevolution of number of OR, TAAR and OlfC genes in ray-finned fishes. (A) OR and TAAR families. (B) OR and OlfC families. (C) TAAR and OlfC families. Coefficient of determination (R^2^), p-value (P) and regression line (solid line) of PGLS analyses are reported. Dashed line shows slope = 1. Dot color code: red, *Polypterus senegalus*; brown *Erpetoichthys calabaricus*; light blue *Polyodon spathula*; dark blue *Acipenser ruthenus*; yellow *Amia calva*; dark green *Atractosteus spatula*; light green *Lepisosteus oculatus*; black (teleost).

Moreover, in most species, the number of OR genes is greater than the number of TAAR or OlfC genes, and often the number of TAAR genes is greater than the number of OlfC genes (**Fig. 3**).

Although the number of ORA genes is less dynamic, particularly in teleosts, species with a high number of OR, TAAR and OlfC genes, such as Polypteriformes or Anguilliformes, tend to have more ORA genes, whereas species with few genes in these three families, such as *Mola mola*, tend to have fewer ORA genes (**Fig. 1)**.

The coevolution of the three dynamic receptor gene families (OR, TAAR, OlfC) is also supported by the correlation between the number of gene losses along the branches of the phylogenetic tree (Pearson’s r = 0.8 between OR and TAAR, 0.52 between OR and OlfC and 0.62 between TAAR and OlfC, all p-values < 2e-16) and gene gains (r = 0.79 between OR and TAAR, 0.78 between OR and OlfC and 0.69 between TAAR and OlfC, all p-values < 2e-16) (**Extended Data Fig. 4**).

The coevolution of the OR, TAAR and OlfC receptor gene families is further supported by a correlation of the number and proportion of pseudogenes, which agrees with similar gene death rates in the three dynamic gene families (**Extended data Fig. 5**).

Together, these results suggest that dramatic changes in evolutionary constraints on the size of the olfactory repertoire occurred several times, with periods of expansion or contraction affecting OR, TAAR and OlfC olfactory receptor families the same way. Hence, they constitute a single evolutionary unit in ray-finned fishes.

### Coevolution of olfactory organ and olfactory gene repertoire

Using data for 72 species of ray-finned fishes (**Supplementary data 1)**, we confirmed the correlation between the number of OR genes and the number of lamellae in the olfactory organ (PGLS; R^2^ = 0.57, p = 1.38e-14, **Fig. 4A**) reported recently for a smaller sample of 35 teleosts and two non-teleost ray-finned fishes ^13^. While no significant correlation was found between the number of lamellae and the number of TAAR genes (PGLS; R^2^ = 0.00177, p = 0.726, **Fig. 4B**), a correlation was found with the number of OlfC genes (PGLS; R^2^ = 0.21, p = 4.55e-05, **Fig. 4C**) and the total number of olfactory receptor genes (PGLS; R^2^ = 0.13, p = 0.00176, **Fig. 4D**).

**Fig. 4.**
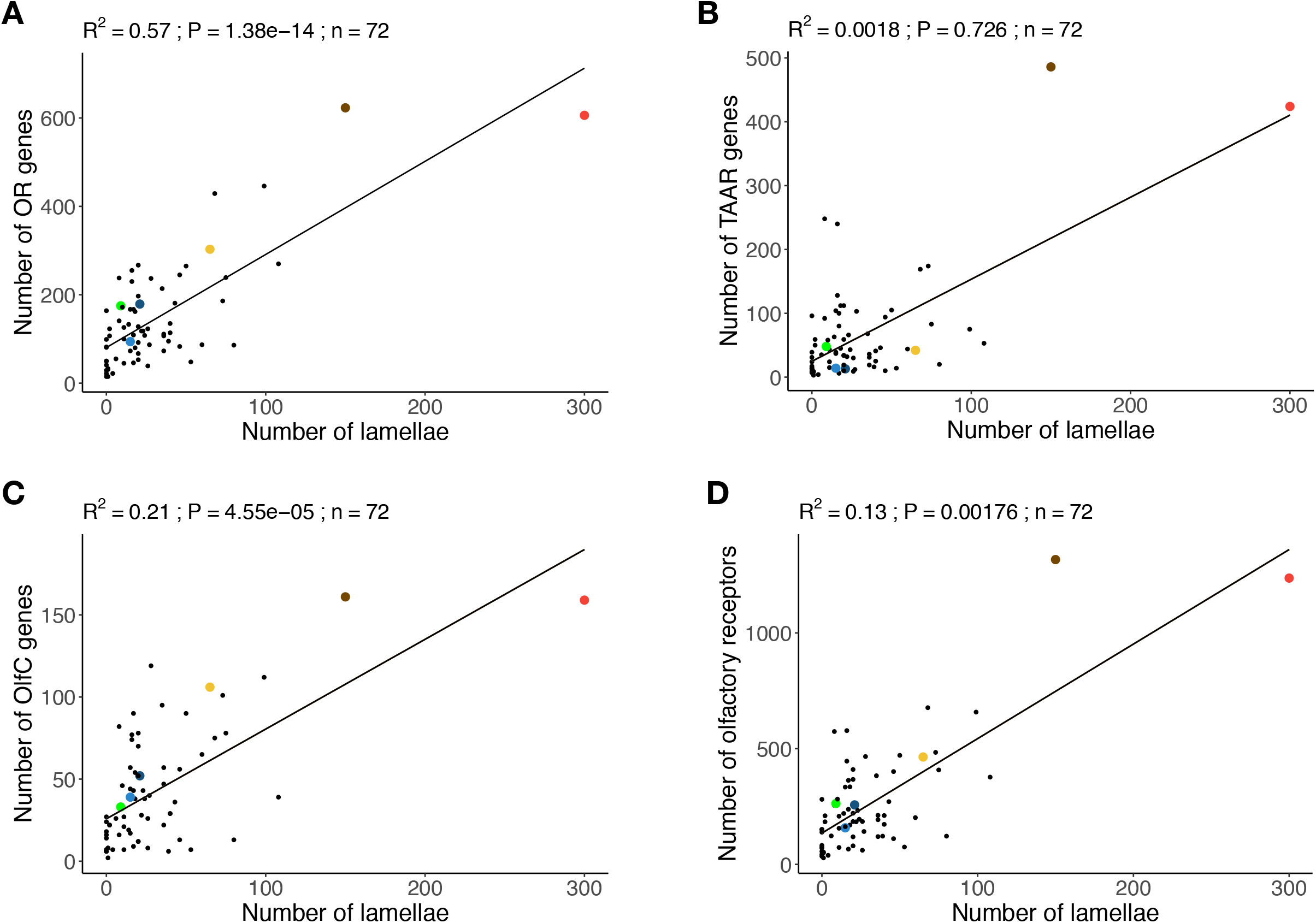
Coevolution of the olfactory gene repertoire and number of olfactory lamellae. (A) OR genes. (B) TAAR genes. (C) OlfC genes. (D) Total olfactory receptor genes. The coefficient of determination (R^2^), the p-value (P) and regression line (solid line) of PGLS analyses are reported. Dot color code: Polypteriformes (red, *Polypterus senegalus*; brown *Erpetoichthys calabaricus*); Acipenseriformes (light blue *Polyodon spathula*; dark blue *Acipenser ruthenus*); Amiiformes (yellow *Amia calva*); Lepisosteiformes (dark green *Atractosteus spatula*; light green *Lepisosteus oculatus*); teleost (black).

The smallest olfactory repertoires occur in Ocean Sunfish *Mola mola* (28 genes) and Broad-nosed Pipefish *Syngnathus typhle* (35 genes). These extreme reductions of olfactory receptor diversity evolved independently and in parallel with the simplification of the olfactory organ, which is a small, flat olfactory epithelium in both species ^13,24^ (**Fig. 2A**). Moreover, *M. mola* has greatly reduced olfactory nerves and reduced olfactory bulbs ^25^.

At the other extreme is the unique organization of the olfactory organ of polypteriforms. In both species studied, the olfactory organ consists of six sectors, each with a rosette-like structure ^14,15^, resulting in many more olfactory lamellae than any other ray-finned fishes (**Fig. 2A**). Polypteriforms also have a much larger olfactory gene repertoire with many more genes in all four gene families than in any other ray-finned fishes (*Polypterus senegalus*: 1237 olfactory receptors, 300 olfactory lamellae; *Erpetoichthys calabaricus:* 1317 olfactory receptors, 150 olfactory lamellae; **Fig. 1**). The two other species studied that had the most olfactory receptor genes also had many olfactory lamellae (*Anguilla anguilla*: 658 olfactory receptors and 99 olfactory lamellae; *Mastacembelus armatus*: 677 olfactory receptors, 68 olfactory lamellae; **Fig. 1**). Interestingly, *P. senegalus, E. calabaricus, A. anguilla* and *M. armatus* are nocturnal (e.g., ^26,27^), perhaps making them more reliant on olfaction. They also have other specializations of the olfactory system, such as prominent, anteriorly directed incurrent narial tubes (**Extended data Fig. 6**). Such tubes direct water flow into the olfactory organ and allow the fish to sample water away from its boundary layer ^28^.

After an extreme contraction of the olfactory gene repertoire and simplification of the olfactory epithelium, a secondary expansion in the gene repertoire occurred in parallel with the reacquisition of a multilamellar epithelium in the tetraodontiform genus *Takifugu*. The genomes of *Takifugu rubripes*, *T. flavidus*, and *T. bimaculatus* have more olfactory genes (156, 124 and 140 respectively) than in other tetraodontiforms with a flat olfactory epithelium, *Dichotomyctere nigroviridis* (70 genes) and *Mola mola* (28 genes). The increased number of genes in the three species of *Takifugu* is due to duplications of OR, TAAR and OlfC genes (**Extended data Fig. 1B-D**). We dissected a specimen of *T. rubripes* and found a non-rosette, but multilamellar, organization of parallel lamellae on the floor of the olfactory chamber that continues on the ventral surface of the nasal bridge between the incurrent and excurrent nares (**Extended data Fig. 7)**. This novel organization supports the hypothesis of a reacquisition of a multilamellar olfactory epithelium in association with secondary expansion of the olfactory receptor gene repertoire.

Together, our results indicate a functional link between receptor diversity and the number of lamellae in the olfactory organ of ray-finned fishes (**Fig. 2B**). In the most extreme cases, this leads to the loss of the rosette (e.g., *Mola mola* and *Syngnathus typhle*) or anatomical innovations with several rosettes (e.g., polypteriforms) or a novel organization of olfactory lamellae (e.g., species of *Takifugu*). This link limits the morpho-genomic space occupied by ray-finned fishes (**Fig. 2B**). Accordingly, we did not observe ray-finned fishes with many olfactory genes and few olfactory lamellae or fishes with few olfactory genes and many olfactory lamellae (**Figs. 2, 4**). Because a large number of olfactory neurons expressing each olfactory receptor is necessary for an efficient olfaction, there is a functional limit to the number of olfactory receptor genes that can be expressed on a given area of olfactory epithelium. This would explain why there are no species with many olfactory receptor genes and few olfactory lamellae. We did not find any examples of macrosmatic fishes with a low number of olfactory receptor genes, which would favor high sensitivity for a small set of odorants. There is probably no functional limit moderating the evolution of such a specialization, and perhaps cartilaginous fishes, which have few olfactory receptor genes and large multilamellar olfactory organs ^29^, may occupy this area of the morpho-genomic space.

## Conclusion

Our analysis of 185 high quality genomes of ray-finned fishes highlights the diversity of the olfactory receptor repertoire. The number of genes is highly dynamic for three (OR, TAAR, OlfC) of the four gene families, but the reasons for large gene gains or losses are still unknown. In marine tetrapods, including cetaceans and sea snakes, extreme reductions in the number of olfactory genes occurred likely because air-adapted olfactory systems were not useful in marine environments ^30^. No such major ecological transition is associated with gene losses of similar magnitude in Syngnathiformes and Tetraodontiformes, and it remains unknown why their olfaction degenerated at both morphological and genomic levels. The complexity of the olfactory organ and large olfactory gene repertoire in Polypteriformes is also surprising. To date, few olfactory receptor genes have been de-orphanized, and such functional information, combined with behavioral studies, may shed light on the dynamics of losses and specializations. Together, our analyses of the olfactory gene repertoire and morphology of the olfactory epithelium show that olfaction is a heterogeneous sensory modality in ray-finned fishes. Our identification of non-model species with particularly poorly developed olfaction (e.g., *Mola mola*) or exceptionally well-developed sense of smell (e.g., *Erpetoichthys calabaricus*) opens new possibilities for comparative and functional research on olfaction.

## Materials and Methods

### Olfactory epithelium data

We surveyed the literature on olfactory organs in fishes and found data on number of lamellae for 60 species for which a draft genome assembly was available. We also dissected the olfactory organ and made lamellae counts for 12 species at the National Museum of Natural History, Washington, DC USA. Literature and specimen data were collected from adults because the number of lamellae often increases with total length (TL); we did not consider sexual dimorphism or individual age, which can impact number of olfactory lamellae ^1,31^. We classified the olfactory epithelium as flat if it had ≤ 2 lamellae and multilamellar if it had > 2 lamellae following Hansen et al. (2005) ^4^. Olfactory lamellae data used in the analyses is summarized in **Supplementary data 1**.

### Genome selection and mining of olfactory receptor genes

Using BUSCO (v5.2.2) ^32^, high quality genomes of 185 ray-finned fishes were selected, including 178 teleost genomes with a BUSCO score > 90% and seven non-teleost genomes with BUSCO score ranging from 81%to 93% (**Supplementary data 1)**.

A time-calibrated phylogenetic tree of ray-finned fishes was downloaded from https://fishtreeoflife.org ^7^ and pruned using the R package ape (v5.0) ^33^ to the 185 species included in our study.

Single-exon genes that code for OR receptors were mined following methods described by Policarpo et al. (2021) ^13^. TAAR, OlfC and ORA genes, which consist of several exons, were identified following Azzouzi et al. (2015) ^34^, with slight modifications. In brief, a TBLASTN ^35^ was performed using known TAAR, OlfC or ORA sequences as queries with a threshold e-value < 1e-10 to select regions containing putative TAAR, OlfC or ORA genes. Non-overlapping hit regions were extracted and extended 5000 bp upstream and downstream using SAMtools ^36^. For each extended non-overlapping hit region, the protein with the best TBLASTN match was aligned to the DNA sequence using EXONERATE (v2.2) ^37^ and the resulting protein-coding sequence was used as query for a BLASTX against a custom database of OR, TAAR, OlfC, ORA and other G protein-coupled receptors (GPCRs). Protein-coding sequences that best matched TAAR, OlfC or ORA receptors were retained and manually curated. Each protein-coding sequence was translated and aligned to known olfactory receptors and other GPCR genes with MAFFT (v7.487) ^38^ and maximum likelihood trees were computed with IQ-TREE (v1.6.12) ^39^. Only protein-coding sequences that clustered with known olfactory receptors by visual inspection using iTOL ^40^ were retained as olfactory receptor genes. When several identical sequences were retrieved in a genome, only one was kept using CD-HIT ^41^.

Retrieved coding sequences were classified as: 1) ‘gene’ if complete and without loss-of-function mutation (premature stop codon or frameshift), 2) ‘pseudogene’ if complete and with at least one loss-of-function mutation, 3) ‘truncated’ if incomplete and without loss-of-function mutation, 4) ‘edge’ if incomplete and less than 30 bp from a contig border.

We assessed the quality of our mining pipeline by comparing the olfactory gene repertoires we identified with those published by other authors for four teleost species. We systematically found more genes than previous studies, in particular, in *P. senegalus*, we identified three times more OR genes (**Supplementary data 2**).

### Phylogenies and gene classification

For each species, we aligned protein sequences coded by putative OR genes with known OR genes ^22^ using MAFFT. A maximum likelihood tree was computed with IQ-TREE and genes were classified according to their position in the tree. To assess the relative diversity of OR subfamilies, a phylogenetic tree with OR genes of 44 species, each species belonging to a different order based on fishtreeoflife (https://fishtreeoflife.org/), was computed. The root was placed between Type I and Type II genes (**Supplementary data 3**). Using MAFFT, putative TAAR genes were aligned with TAARs and non-TAAR GPCRs genes obtained from Dieris et al. (2021) ^42^. A maximum likelihood tree was computed with IQ-TREE and genes were classified according to their position in the phylogenetic tree (**Supplementary data 3**). The same method was used for putative OlfC and ORA genes. For putative OlfC genes, we used genes from Yang et al. (2019) ^19^ and CasR and V2R2 genes as outgroups (**Supplementary data 3**). For ORA sequences, we used genes from Zapilko et al. (2016) ^18^ and T2R genes as an outgroup (**Supplementary data 3**).

Pseudogenes, truncated genes and edge gene classification was based on the best blastx match.

### Phylogenetic generalized least-squares regression analyses

We estimated phylogenetic signal (Pagel’s λ) of each trait with the function phylosig in the R package phytools with the option test = TRUE ^43^. The R package caper (v1.0.1) ^44^ was used to perform phylogenetic generalized least square analyses using the function “pgls” with lambda = “ML” (**Supplementary data 1**).

### Gene tree – Species tree reconciliation

The number of gene gains and number of gene losses along each branch of the species phylogenetic tree were inferred using the gene tree – species tree reconciliation method. The OR family is large, as described previously ^13^, and thus OR genes belonging to different subfamilies were aligned separately. For the smaller TAAR, OlfC and ORA gene families, one alignment was obtained for each gene family separately. All alignments were obtained using MAFFT. Maximum likelihood trees were computed with IQ-TREE. Nodes with low bootstrap values (< 90%) were collapsed into polytomies using the R package ape. We then used Treerecs ^23^ to root and reconcile genes trees with the species tree.

For each olfactory receptor family, we computed birth and death rates using equations in Niimura et al. (2014) ^45^ excluding branches with length < 2 Mya because differences in gene retrieval and genome qualities greatly impacted inferred birth and death rate ^13^.

## Data availability

All sequences extracted in this study are available on https://figshare.com/articles/dataset/Olfactory_receptor_sequences_for_185_ray-finned_fishes/17061632.

## Supplementary Material

Supplementary data 1. Sheet 1: NCBI Assembly accession and assembly level of the 185 genomes studied, their species name in NCBI and in Eschmeyer’s Catalog of Fishes. Sheet 2: results of BUSCO analyses on the 185 genomes studied. Sheet 3: species’ name and order based on the taxonomy of fishtreeoflife.org. Sheet 4: summary of the number of genes in each olfactory receptor family for every species. Sheet 5: Olfactory epithelium shape and number of lamellae in 72 ray-finned fishes for which a genome assembly is available. Sheet 6: phylogenetic signal of the number of gene in each family and of the number of lamellae in the epithelium computed with phytools. Values of phylogenetic regression described in this study are also given.

Supplementary data 2. Comparison of olfactory receptor gene repertoires from the present and previous studies. (A) Summary of the number of TAAR genes retrieved in our study and previous studies of four teleost species. (B) Summary of the number of OlfC genes retrieved in our study and previous studies of four teleost species. (C) Summary of the number of ORA genes retrieved in our study and previous studies of four teleost species. (D) Phylogenetic tree of *Danio rerio* TAAR genes retrieved in Hashiguchi and Nishida 2007 and our study. (E) Phylogenetic tree of *Gasterosteus aculeatus* TAAR genes retrieved in Azzouzi et al. 2015 and our study. (F) Phylogenetic tree of *Oryzias latipes* TAAR genes retrieved in Azzouzi et al. 2015 and our study. (G) Phylogenetic tree of *Takifugu rubripes* TAAR genes retrieved in Hashiguchi and Nishida 2007 and our study. (H) Phylogenetic tree of *Danio rerio* OlfC genes retrieved in Yang et al. 2019 and our study. (I) Phylogenetic tree of *Gasterosteus aculeatus* OlfC genes retrieved in Yang et al. 2019 and our study. (J) Phylogenetic tree of *Oryzias latipes* OlfC genes retrieved in Yang et al. 2019 and our study. (K) Phylogenetic tree of *Takifugu rubripes* OlfC genes retrieved in Yang et al. 2019 and our study. (L) Phylogenetic tree of *Danio rerio* ORA genes retrieved in Zapilko and Korsching 2016 and our study. (M) Phylogenetic tree of *Gasterosteus aculeatus* ORA genes retrieved in Zapilko and Korsching 2016 and our study. (N) Phylogenetic tree of *Oryzias latipes* ORA genes retrieved in Zapilko and Korsching 2016 and our study. (O) Phylogenetic tree of *Takifugu rubripes* ORA genes retrieved in Zapilko and Korsching 2016 and our study. (P) Phylogenetic tree of *Polypterus senegalus* OR genes retrieved in Bi X et al. 2021 and our study.

Supplementary data 3. (A) Phylogeny of OR genes from 44 species representing 44 orders of ray-finned fishes sampled in this study. Branches are colored according to the gene subfamily classification. (B) Phylogeny of all TAAR genes retrieved from 185 ray-finned fishes. Branches are colored according to gene family classification. Outgroup sequences (nonTAAR GPCRs) are colored in black. (C) Phylogeny of all OlfC genes retrieved from 185 ray-finned fish. Branches are colored according to the gene subfamily classification. Outgroup sequences (CasR and V2R2 genes) are colored in black. (D) Phylogeny of all ORA genes retrieved from 185 ray-finned fish. Branches are colored according to the gene subfamily classification. Outgroup sequences (T2R genes) are colored in black.

## Acknowledgments

This work was supported by a collaborative grant from Institut Diversité Ecologie et Evolution du Vivant (to S.R. and D.C).

M.P. was supported by a PhD fellowship from the French Ministry of Research.

J. Galbraith provided the examined specimen of *Mola mola*.

## Author Contributions

M.P., K.B., S.R and D.C. conceptualized the project. M.P. carried out phylogenetic and statistical analyses. K.B. obtained morphological data. P.L. produced fish drawings. P.L., L.L., J.-C.S. and S.R. contributed to data analyses. D.C. supervised phylogenetic and statistical analyses. M.P. and K.B made the figures. D.C., M.P. and K.B. wrote the manuscript. All authors commented on the manuscript and agreed to its final version.

## Legends Extended data

**Extended data Fig. 1.**
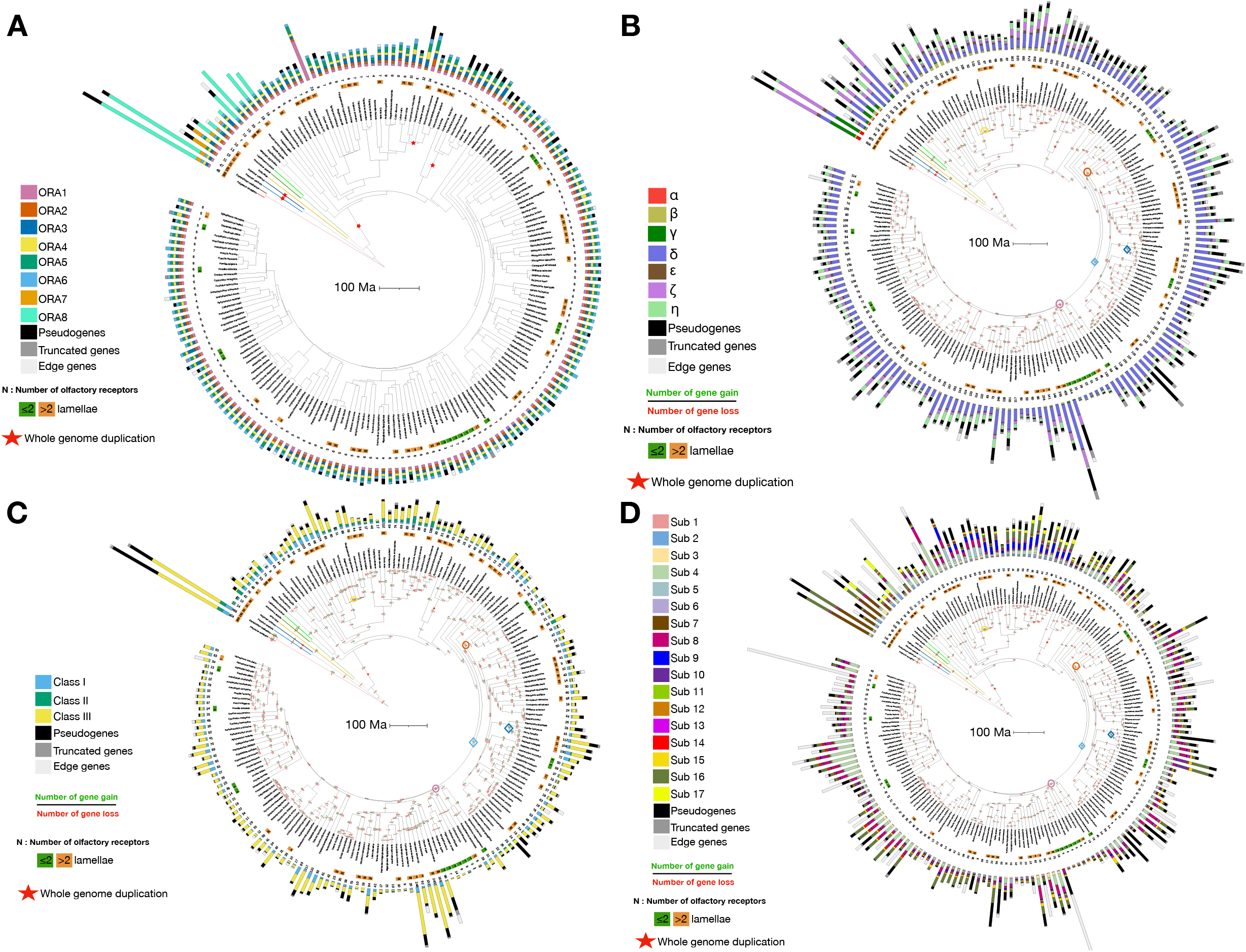
Diversity of the olfactory receptor gene repertoire in ray-finned fishes. Time-calibrated phylogeny from https://fishtreeoflife.org/. (A) For each species, a multiple values barplot represents the number of functional genes, pseudogenes, truncated and edge genes in (A) ORA family, (B) OR family, (C) TAAR family, (D) OlfC family. When available, the olfactory epithelium shape and the number of lamellae is indicated. Inferred numbers of gene gains and losses are provided on branch of the trees. Whole-genome duplications are indicated by red stars. The branches associated with the three highest death rates and the two highest birth rates of the OR, TAAR and OlfC families are indicated by circles and diamonds, respectively, with color code as in Fig. 1. The trees were annotated and visualized using iTOL.

**Extended data Fig. 2.**
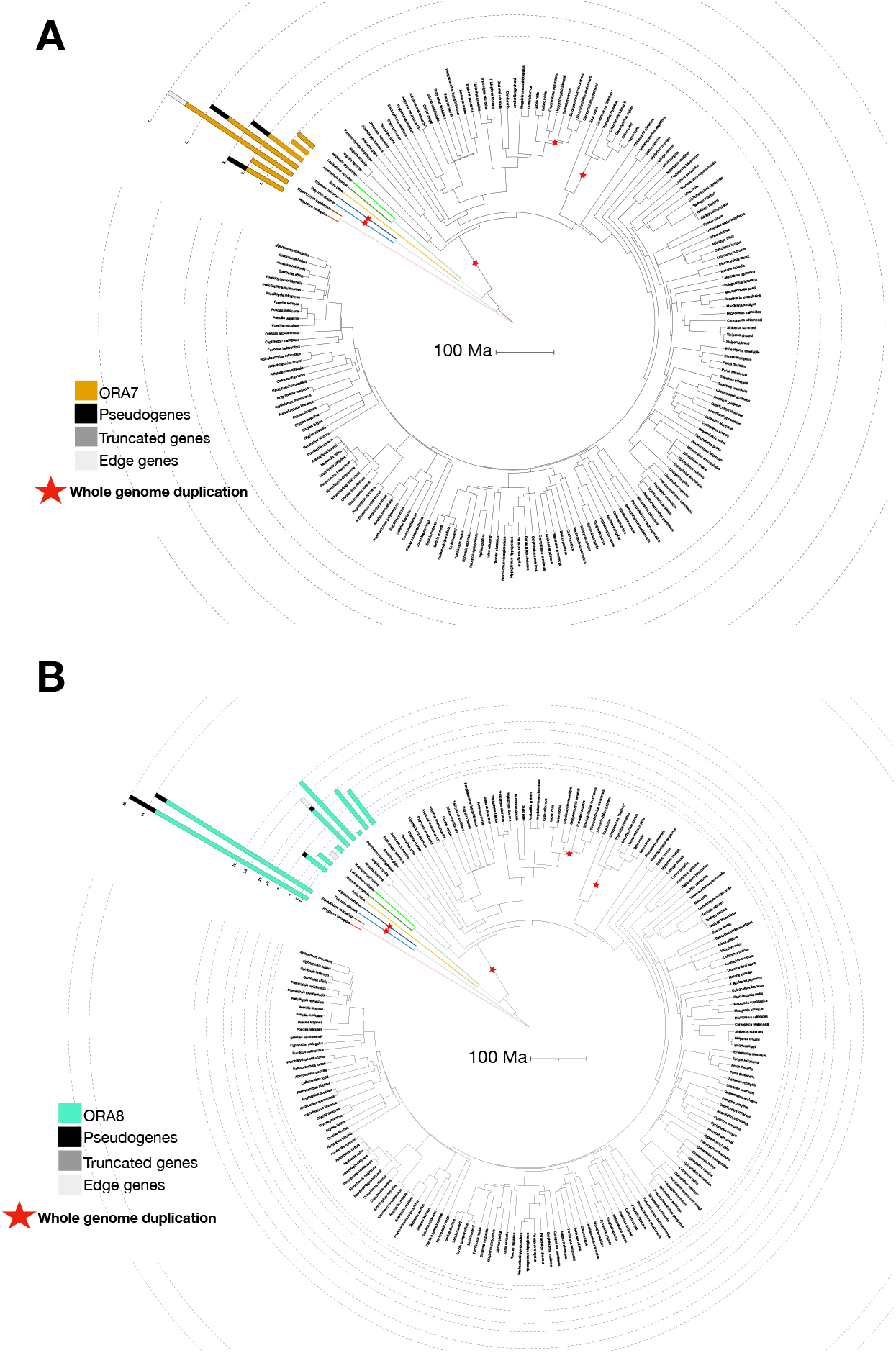
Distribution of ORA7 and ORA8 subfamilies in ray-finned fishes. Time-calibrated phylogeny from https://fishtreeoflife.org/. (A) ORA7, (B) ORA8.

**Extended data Fig. 3.**
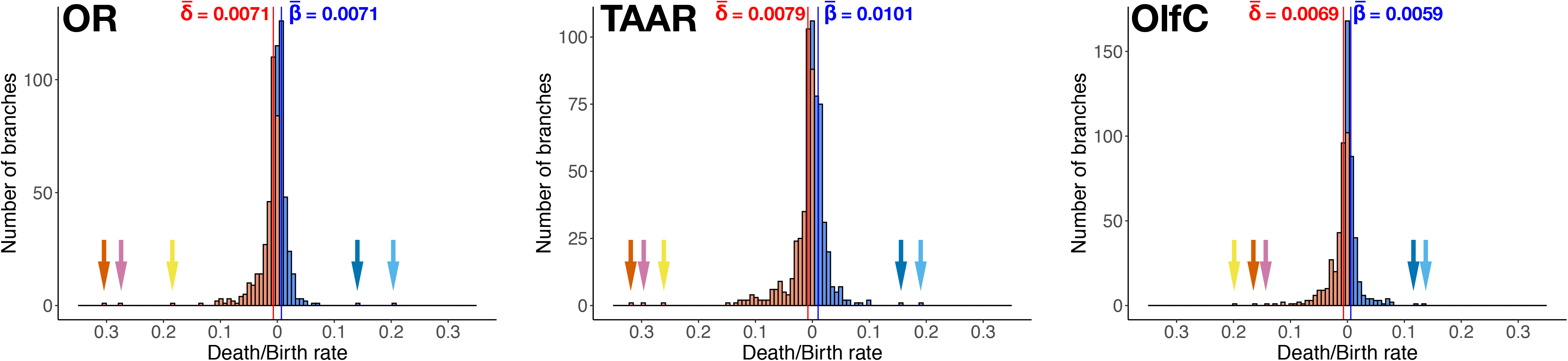
Distribution of birth and death rates of OR, TAAR, and OlfC genes in ray-finned fishes. 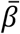: mean birth rate; 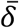: mean death rate. The two highest birth rates and the three highest death rates are indicated by colored arrows. Arrow colors correspond to the colors of circles and diamonds in Fig. 1 showing branches with high birth and death rates.

**Extended data Fig. 4.**
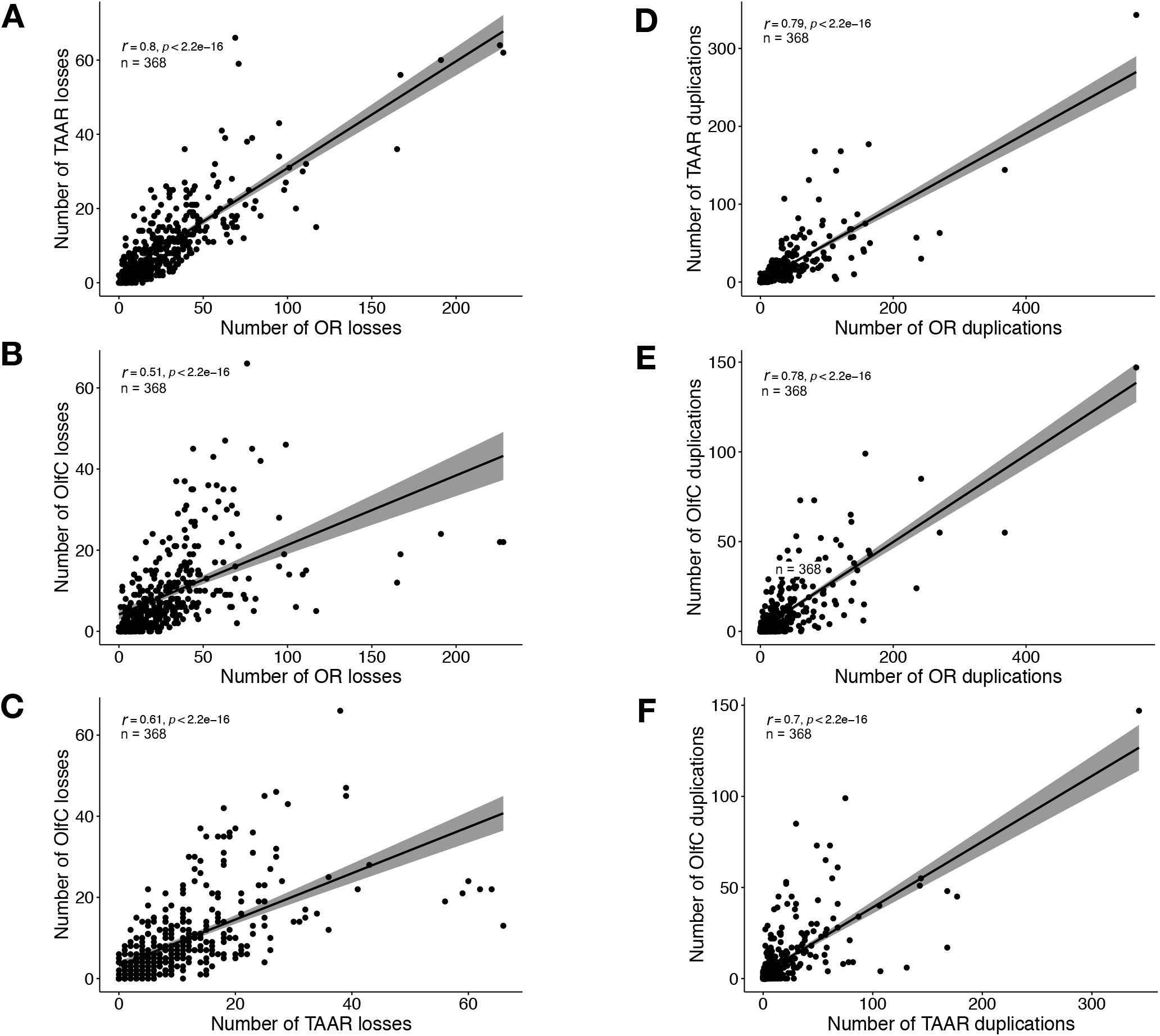
Correlation of the number of gene losses (or gene gains) between gene families, estimated using the 368 branches of the phylogenetic tree. (A,D) Correlations between OR and TAAR families; (B,E) correlations between OR and OlfC families; (C,F) correlations between TAAR and OlfC families.

**Extended data Fig. 5.**
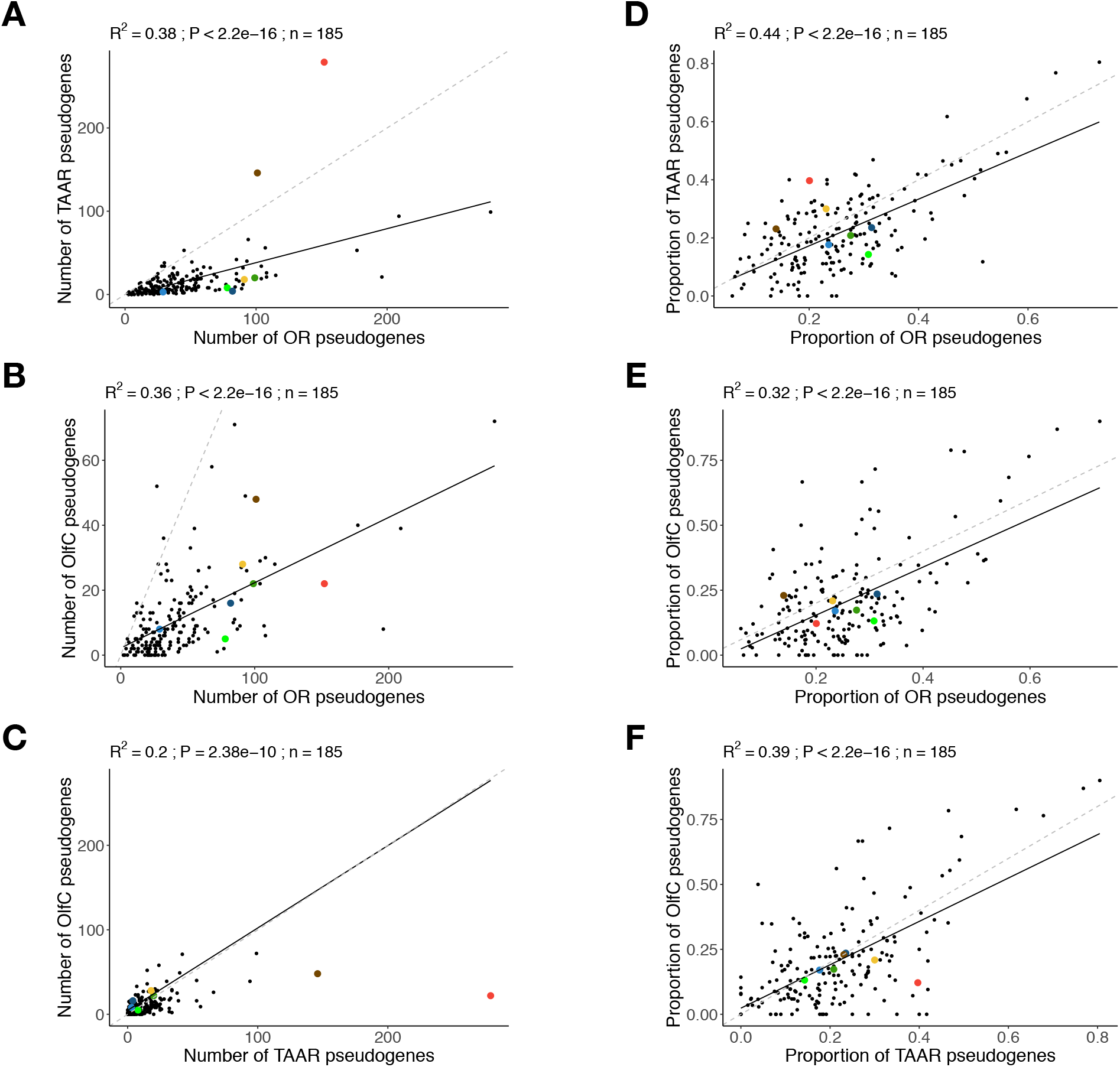
Correlation between the number of OR, TAAR and OlfC pseudogenes in 185 ray-finned fishes. (A) Phylogenetic generalized linear regression (Pagel’s λ model) between the number of OR and TAAR pseudogenes, (B) OR and OlfC pseudogenes, (C) TAAR and OlfC pseudogenes. (D) Phylogenetic generalized linear regression (Pagel’s λ model) between the proportion of OR and TAAR pseudogenes, (E) OR and OlfC pseudogenes, (F) TAAR and OlfC pseudogenes. The coefficient of determination (R2), the p-value (P) and the regression line (solid line) of PGLS analyses are reported. Dotted line: slope = 1. Dot color code: as in Fig. 1.

**Extended data Fig. 6.**
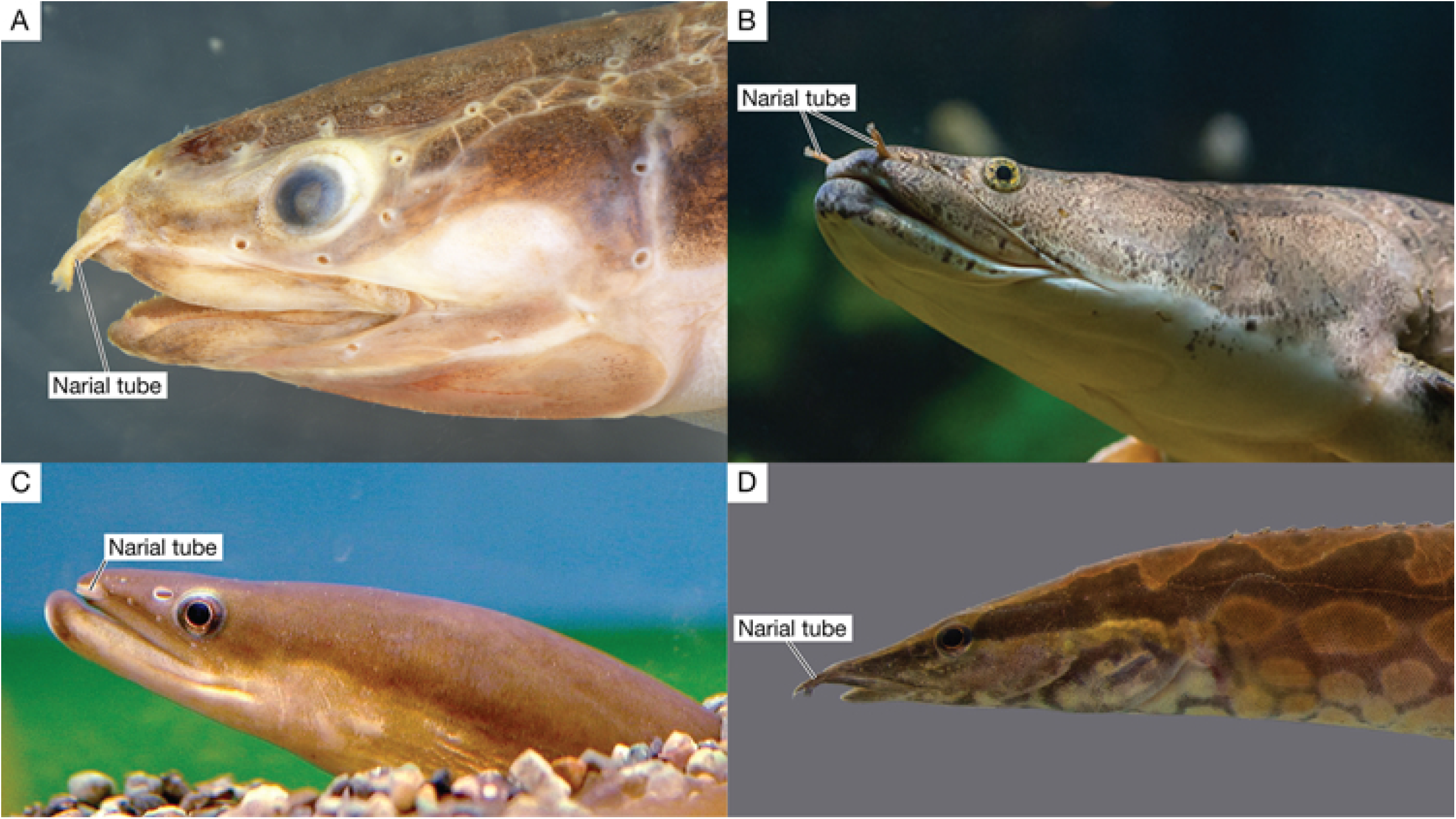
Narial tubes of four species of ray-finned fishes with complex olfactory organs and large gene repertoire. (A) *Erpetoichthys calabaricus*; photograph by Katherine E. Bemis, NOAA National Systematics Lab. (B) *Polypterus senegalus*, photograph by Basal Zoo. (C) Representative Anguillidae, *Anguilla japonica*, photograph by unknown. (D)*Mastacembelus armatus*, photograph by Zach Randall, Florida Museum of Natural History.

**Extended data Fig. 7.**
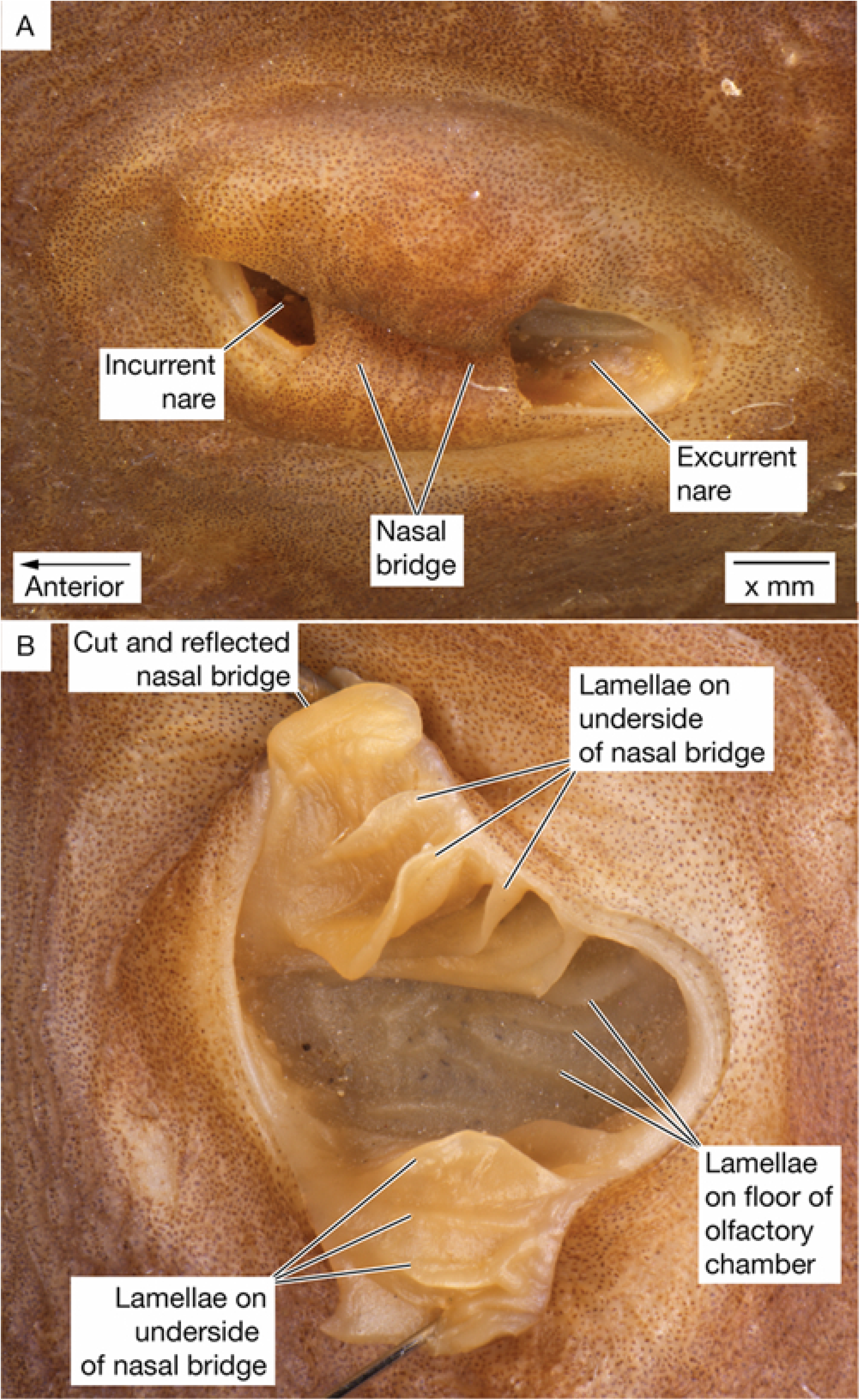
*Takifugu rubripes*, USNM 57620, 290 mm TL. (A) Bridge of tissue between excurrent and incurrent nares. (B) Bridge of tissue cut and reflected to show lamellae in olfactory organ. Note that lamellae are not arranged in a rosette, but parallel to each other in a circle under the bridge and over the floor of the olfactory organ.

