## Supplementary data 2 for "Coevolution of the olfactory organ and its receptor repertoire in ray-finned fishes"

A

| TAAR pipeline assessment | Present study | | | Previous studies | | | Reference |
|---|---|---|---|---|---|---|---|
| | F | I | P | F | I | P | |
| *Danio rerio* | 112 | 1 | 9 | 109 | N/A | 10 | Hashiguchi and Nishida 2007 |
| *Gasterosteus aculeatus* | 50 | 2 | 15 | 50 | N/A | 15 | Azzouzi et al. 2015 |
| *Oryzias latipes* | 39 | 2 | 4 | 27 | N/A | 7 | Azzouzi et al. 2015 |
| *Takifugu rubripes* | 24 | 0 | 3 | 13 | N/A | 6 | Hashiguchi and Nishida 2007 |

B

| OlfC pipeline assessment | Present study | | | Previous studies | | | Reference |
|---|---|---|---|---|---|---|---|
| | F | I | P | F | I | P | |
| *Danio rerio* | 54 | 3 | 2 | 53 | 1 | 1 | Yang et al. 2019 |
| *Gasterosteus aculeatus* | 22 | 2 | 2 | 13 | 1 | 1 | Yang et al. 2019 |
| *Oryzias latipes* | 24 | 0 | 2 | 16 | 1 | 2 | Yang et al. 2019 |
| *Takifugu rubripes* | 27 | 0 | 4 | 16 | 2 | 3 | Yang et al. 2019 |

**C**

| ORA pipeline assessment | Present study | | | Previous studies | | | Reference |
|---|---|---|---|---|---|---|---|
| | F | I | P | F | I | P | |
| *Danio rerio* | 7 | 0 | 0 | 7 | N/A | N/A | Zapilko and Korsching 2016 |
| *Gasterosteus aculeatus* | 5 | 0 | 1 | 6 | N/A | N/A | Zapilko and Korsching 2016 |
| *Oryzias latipes* | 7 | 0 | 1 | 7 | N/A | N/A | Zapilko and Korsching 2016 |
| *Takifugu rubripes* | 5 | 0 | 1 | 5 | N/A | N/A | Zapilko and Korsching 2016 |

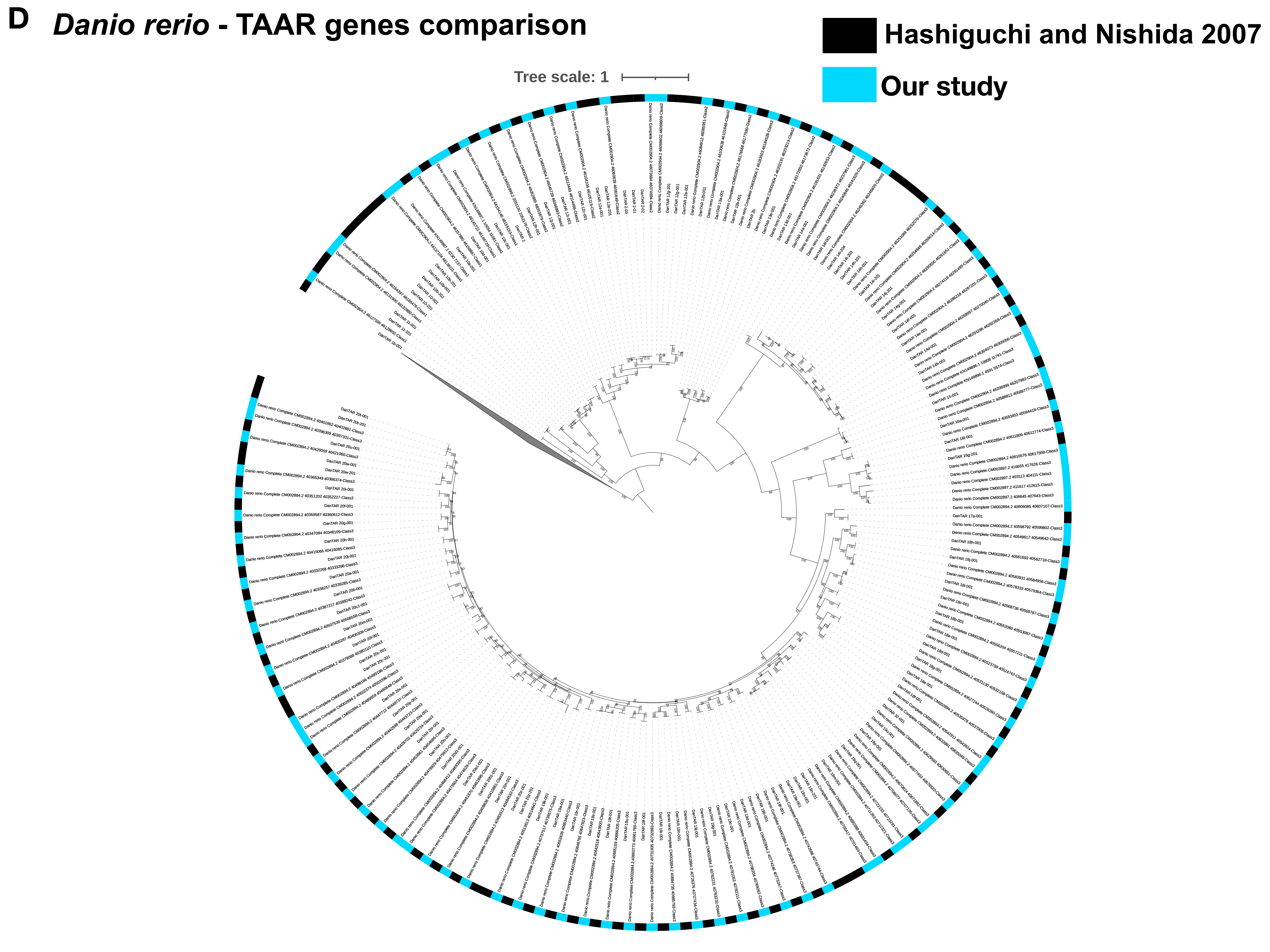
D
Danio rerio - TAAR genes comparison
Hashiguchi and Nishida 2007
Our study
Tree scale: 1

### E *Gasterosteus aculeatus* - TAAR genes comparison

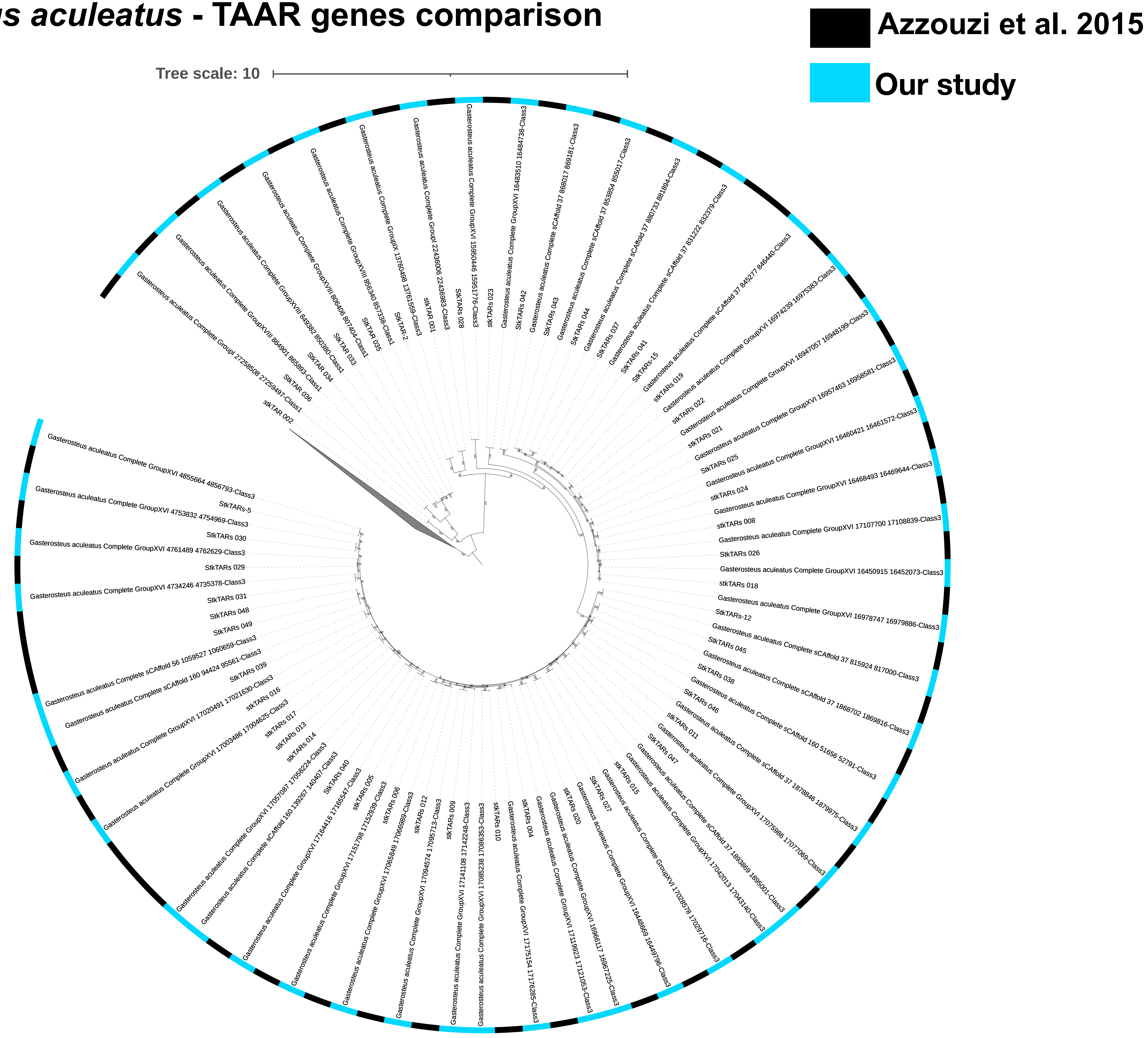

F *Oryzias latipes* - TAAR genes comparison

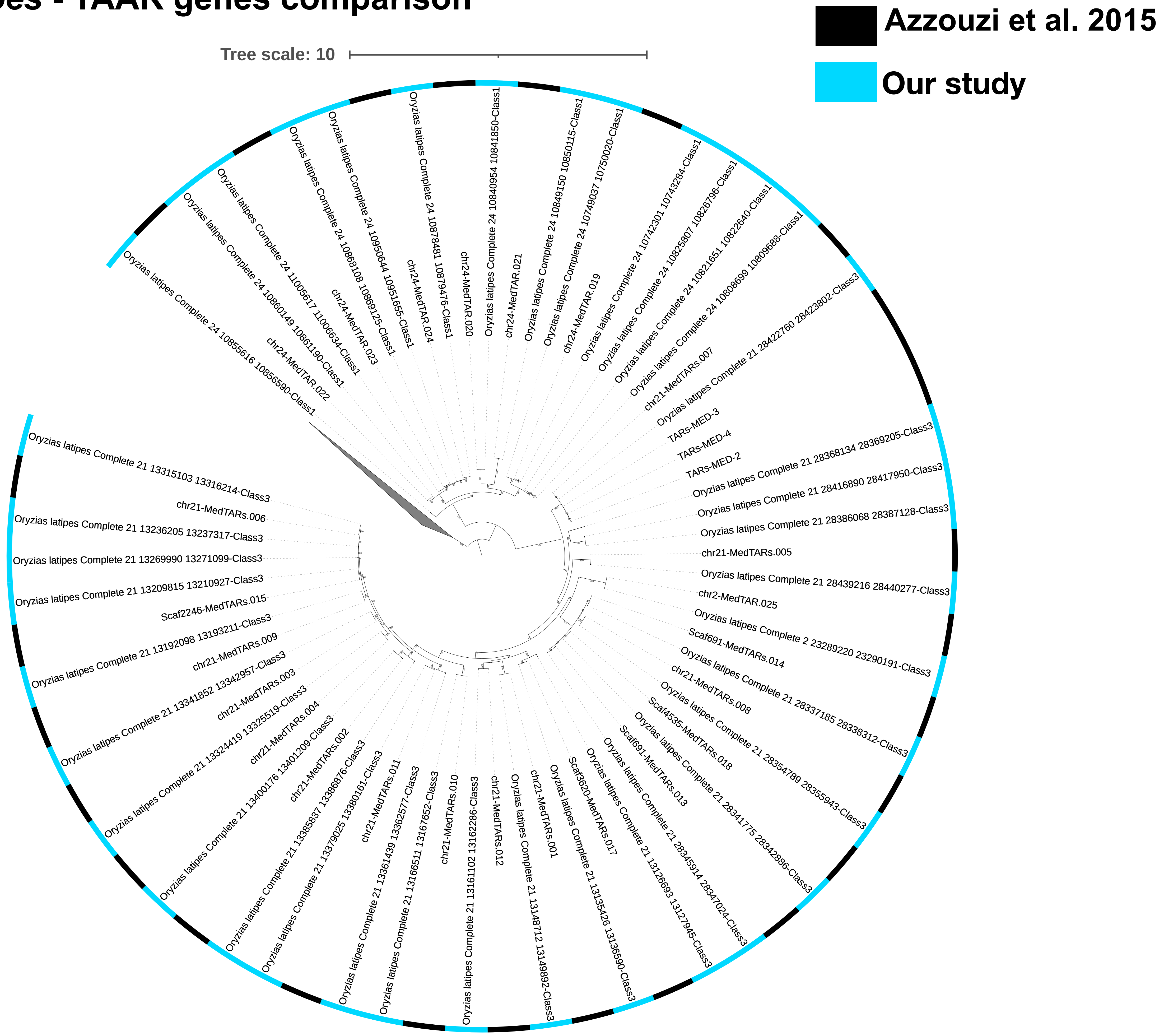

G *Takifugu rubripes* - TAAR genes comparison

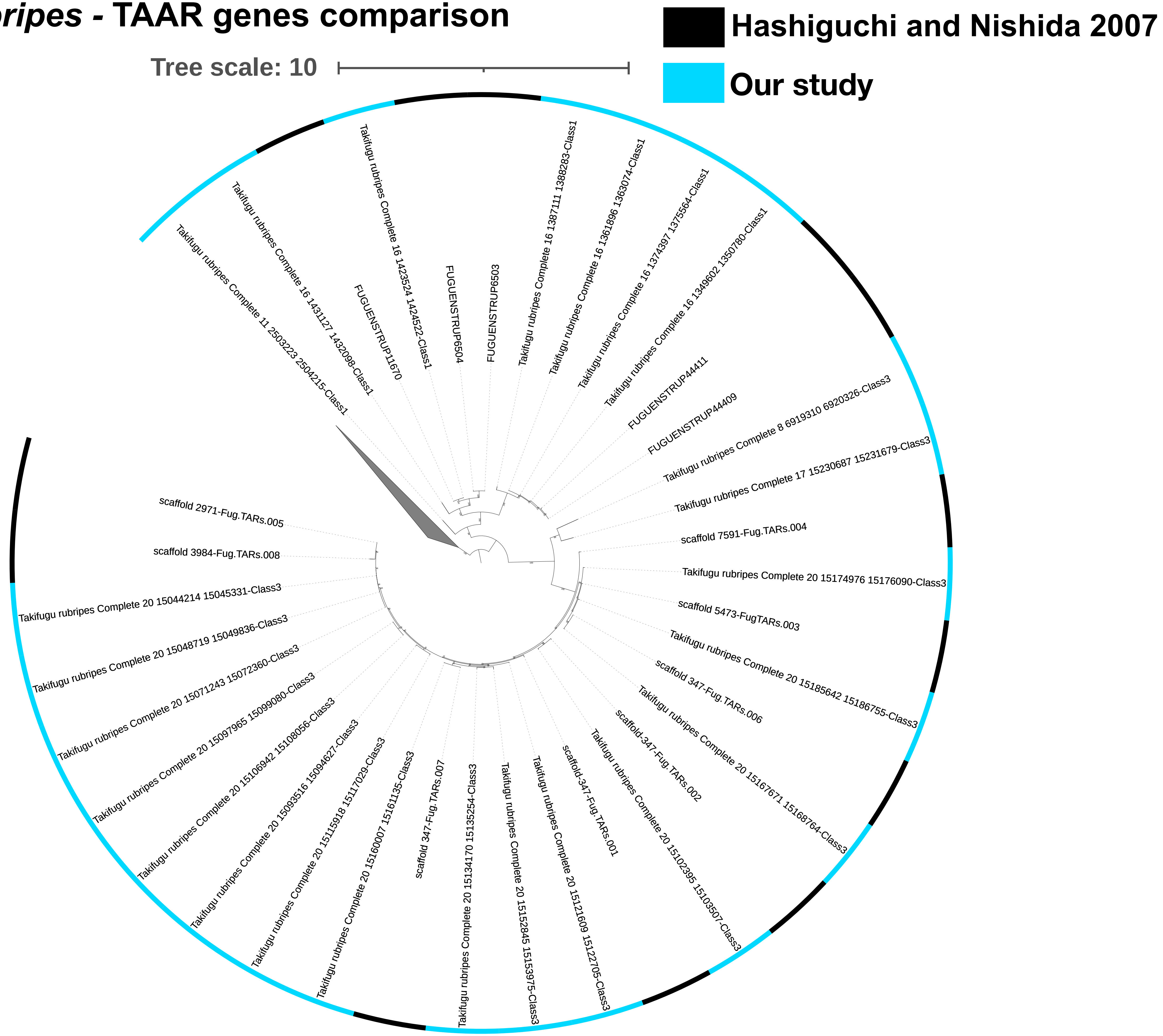

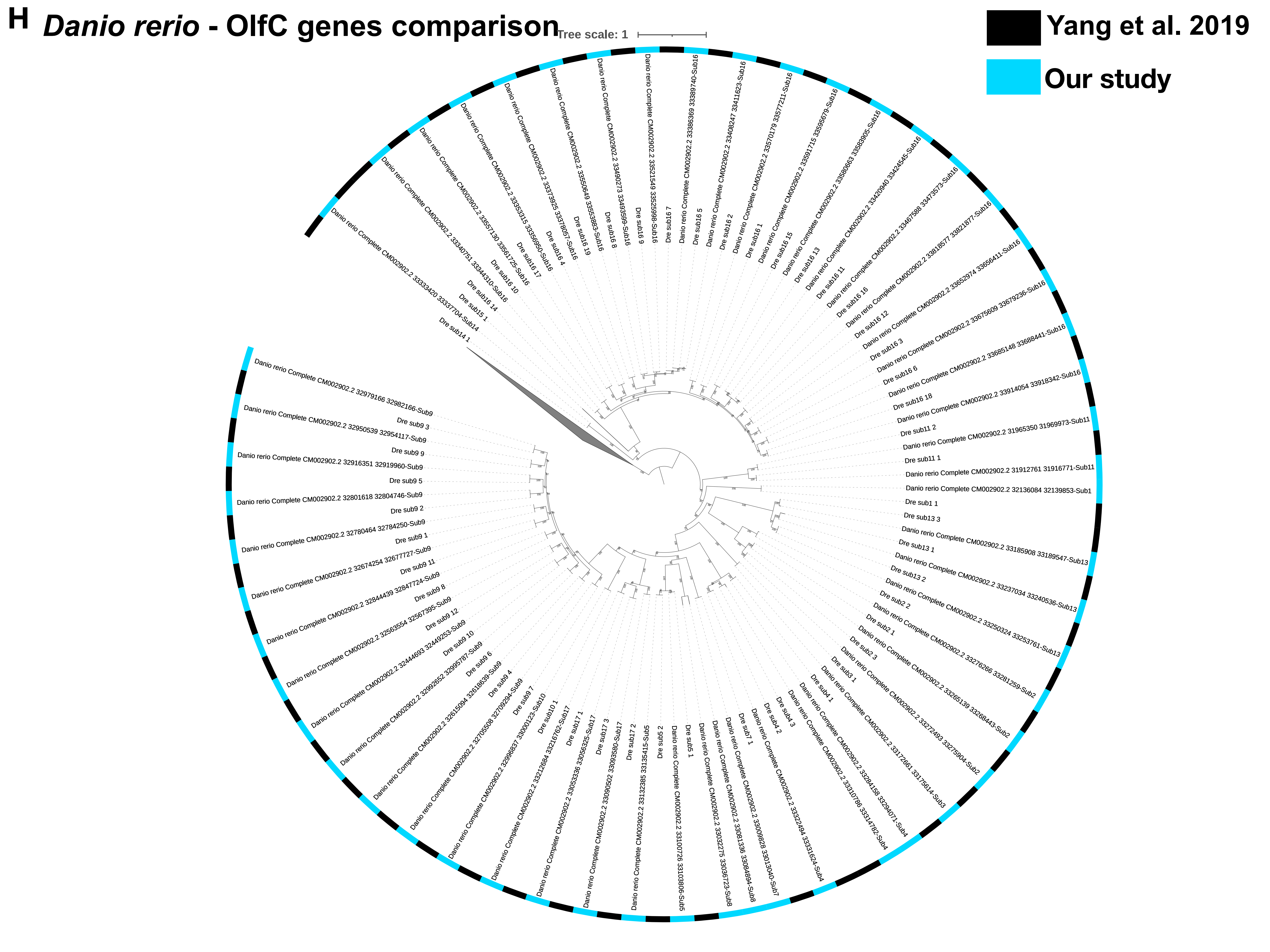
H
Danio rerio - OlfC genes comparison
Tree scale: 1
Yang et al. 2019
Our study
Dre sub14 1
Danio rerio Complete CM002902.2 33333420 33337704-Sub14
Dre sub15 1
Dre sub16 14
Danio rerio Complete CM002902.2 33340751 33344310-Sub16
Dre sub16 10
Danio rerio Complete CM002902.2 33557130 33561725-Sub16
Dre sub16 17
Danio rerio Complete CM002902.2 33353315 33356950-Sub16
Dre sub16 4
Danio rerio Complete CM002902.2 33373925 33378057-Sub16
Dre sub16 19
Danio rerio Complete CM002902.2 33550649 33553883-Sub16
Dre sub16 8
Danio rerio Complete CM002902.2 33490273 33493599-Sub16
Dre sub16 9
Danio rerio Complete CM002902.2 33521549 33525998-Sub16
Dre sub16 7
Danio rerio Complete CM002902.2 33386369 33389740-Sub16
Dre sub16 5
Dre sub16 2
Danio rerio Complete CM002902.2 33408247 33411623-Sub16
Dre sub16 1
Danio rerio Complete CM002902.2 33570179 33577211-Sub16
Dre sub16 15
Danio rerio Complete CM002902.2 33591715 33595679-Sub16
Dre sub16 13
Danio rerio Complete CM002902.2 33580663 33583905-Sub16
Dre sub16 11
Danio rerio Complete CM002902.2 33420940 33424545-Sub16
Dre sub16 16
Danio rerio Complete CM002902.2 33467588 33473573-Sub16
Dre sub16 12
Danio rerio Complete CM002902.2 33818577 33821877-Sub16
Dre sub16 3
Danio rerio Complete CM002902.2 33652974 33656411-Sub16
Dre sub16 6
Danio rerio Complete CM002902.2 33675609 33679236-Sub16
Dre sub16 18
Danio rerio Complete CM002902.2 33685148 33688441-Sub16
Danio rerio Complete CM002902.2 33914054 33918342-Sub16
Dre sub11 2
Danio rerio Complete CM002902.2 31965350 31969973-Sub11
Dre sub11 1
Danio rerio Complete CM002902.2 31912761 31916771-Sub11
Danio rerio Complete CM002902.2 32136084 32139853-Sub1
Dre sub1 1
Dre sub13 3
Danio rerio Complete CM002902.2 33185908 33189547-Sub13
Dre sub13 1
Danio rerio Complete CM002902.2 33237034 33240536-Sub13
Dre sub13 2
Danio rerio Complete CM002902.2 33250324 33253761-Sub13
Dre sub2 2
Danio rerio Complete CM002902.2 33276266 33281259-Sub2
Dre sub2 1
Danio rerio Complete CM002902.2 33265139 33268443-Sub2
Dre sub2 3
Danio rerio Complete CM002902.2 33272493 33275904-Sub2
Dre sub3 1
Danio rerio Complete CM002902.2 33172661 33175614-Sub3
Dre sub4 1
Danio rerio Complete CM002902.2 33284158 33294071-Sub4
Danio rerio Complete CM002902.2 33310786 33314782-Sub4
Dre sub4 3
Danio rerio Complete CM002902.2 33322494 33331624-Sub4
Dre sub4 2
Danio rerio Complete CM002902.2 33009828 33013040-Sub7
Dre sub7 1
Danio rerio Complete CM002902.2 33081336 33084894-Sub8
Danio rerio Complete CM002902.2 33032275 33036723-Sub8
Dre sub5 1
Danio rerio Complete CM002902.2 33100726 33103808-Sub5
Dre sub5 2
Danio rerio Complete CM002902.2 33132385 33135415-Sub5
Dre sub17 2
Danio rerio Complete CM002902.2 33090502 33093580-Sub17
Dre sub17 3
Danio rerio Complete CM002902.2 33053336 33056325-Sub17
Dre sub17 1
Danio rerio Complete CM002902.2 33212684 33216762-Sub17
Dre sub10 1
Danio rerio Complete CM002902.2 32996837 33000123-Sub10
Dre sub9 7
Danio rerio Complete CM002902.2 32705508 32709294-Sub9
Dre sub9 4
Danio rerio Complete CM002902.2 32615094 32618639-Sub9
Dre sub9 6
Danio rerio Complete CM002902.2 32992652 32995787-Sub9
Dre sub9 10
Danio rerio Complete CM002902.2 32444693 32449253-Sub9
Dre sub9 12
Danio rerio Complete CM002902.2 32563554 32567395-Sub9
Dre sub9 8
Danio rerio Complete CM002902.2 32844439 32847724-Sub9
Dre sub9 11
Danio rerio Complete CM002902.2 32674254 32677727-Sub9
Dre sub9 1
Danio rerio Complete CM002902.2 32780464 32784250-Sub9
Dre sub9 2
Danio rerio Complete CM002902.2 32801618 32804746-Sub9
Dre sub9 5
Danio rerio Complete CM002902.2 32916351 32919960-Sub9
Dre sub9 9
Danio rerio Complete CM002902.2 32950539 32954117-Sub9
Dre sub9 3
Danio rerio Complete CM002902.2 32979166 32982166-Sub9

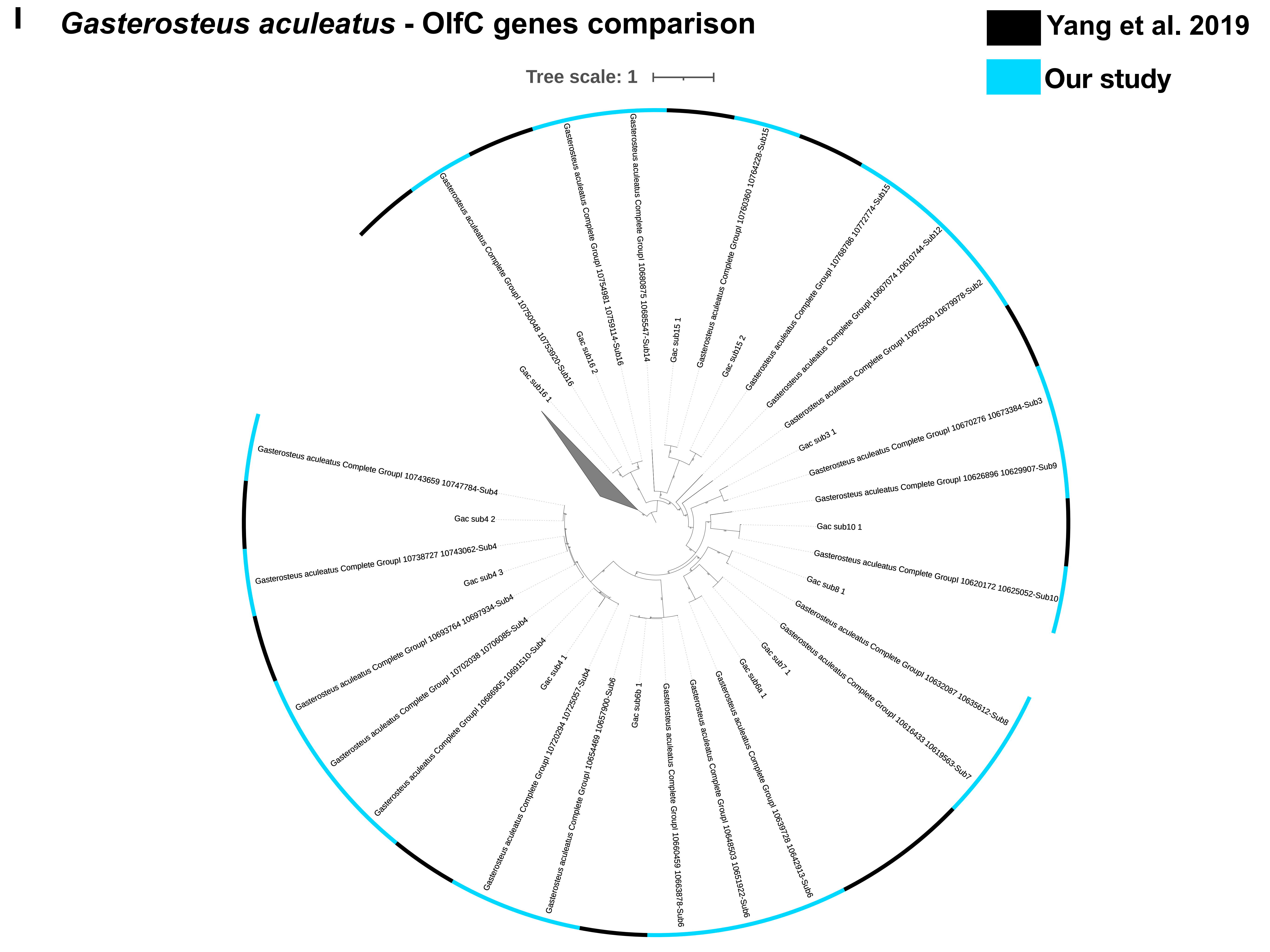
I
Gasterosteus aculeatus - OlfC genes comparison
Yang et al. 2019
Our study
Tree scale: 1
Gac sub16 1
Gasterosteus aculeatus Complete GroupI 10750048 10753920-Sub16
Gac sub16 2
Gasterosteus aculeatus Complete GroupI 10754981 10759114-Sub16
Gasterosteus aculeatus Complete GroupI 10680875 10685547-Sub14
Gac sub15 1
Gasterosteus aculeatus Complete GroupI 10760360 10764228-Sub15
Gac sub15 2
Gasterosteus aculeatus Complete GroupI 10768786 10772774-Sub15
Gasterosteus aculeatus Complete GroupI 10607074 10610744-Sub12
Gasterosteus aculeatus Complete GroupI 10675500 10679978-Sub2
Gac sub3 1
Gasterosteus aculeatus Complete GroupI 10670276 10673384-Sub3
Gasterosteus aculeatus Complete GroupI 10626896 10629907-Sub9
Gac sub10 1
Gasterosteus aculeatus Complete GroupI 10620172 10625052-Sub10
Gac sub8 1
Gasterosteus aculeatus Complete GroupI 10632087 10635612-Sub8
Gasterosteus aculeatus Complete GroupI 10616433 10619563-Sub7
Gac sub7 1
Gac sub6a 1
Gasterosteus aculeatus Complete GroupI 10639728 10642913-Sub6
Gasterosteus aculeatus Complete GroupI 10648503 10651922-Sub6
Gasterosteus aculeatus Complete GroupI 10660459 10663878-Sub6
Gac sub6 1
Gasterosteus aculeatus Complete GroupI 10654469 10657900-Sub6
Gasterosteus aculeatus Complete GroupI 10720294 10725057-Sub4
Gac sub4 1
Gasterosteus aculeatus Complete GroupI 10686905 10691510-Sub4
Gasterosteus aculeatus Complete GroupI 10702038 10706085-Sub4
Gasterosteus aculeatus Complete GroupI 10693764 10697934-Sub4
Gac sub4 3
Gasterosteus aculeatus Complete GroupI 10738727 10743062-Sub4
Gac sub4 2
Gasterosteus aculeatus Complete GroupI 10743659 10747784-Sub4

J *Oryzias latipes* - OlfC genes comparison

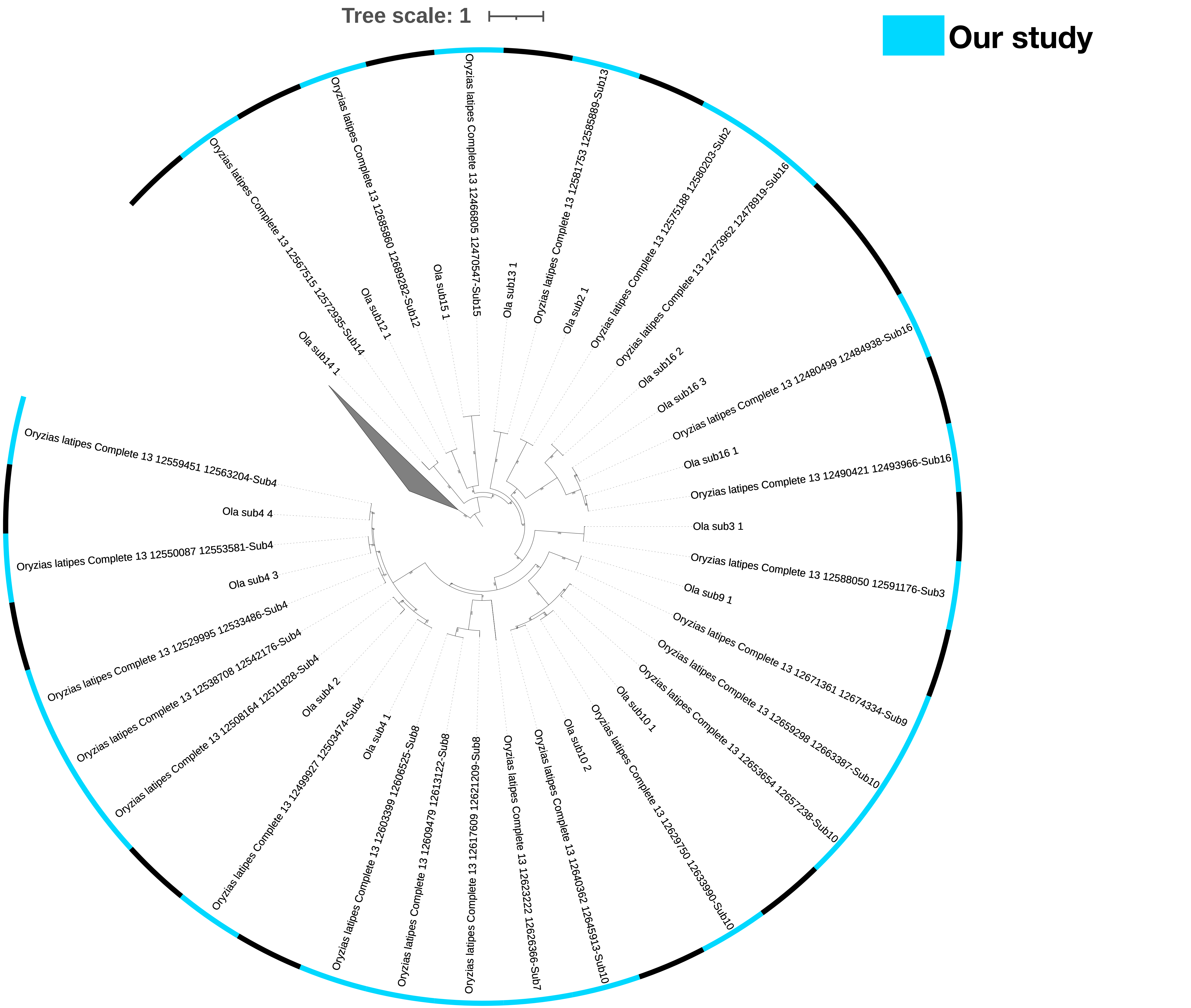

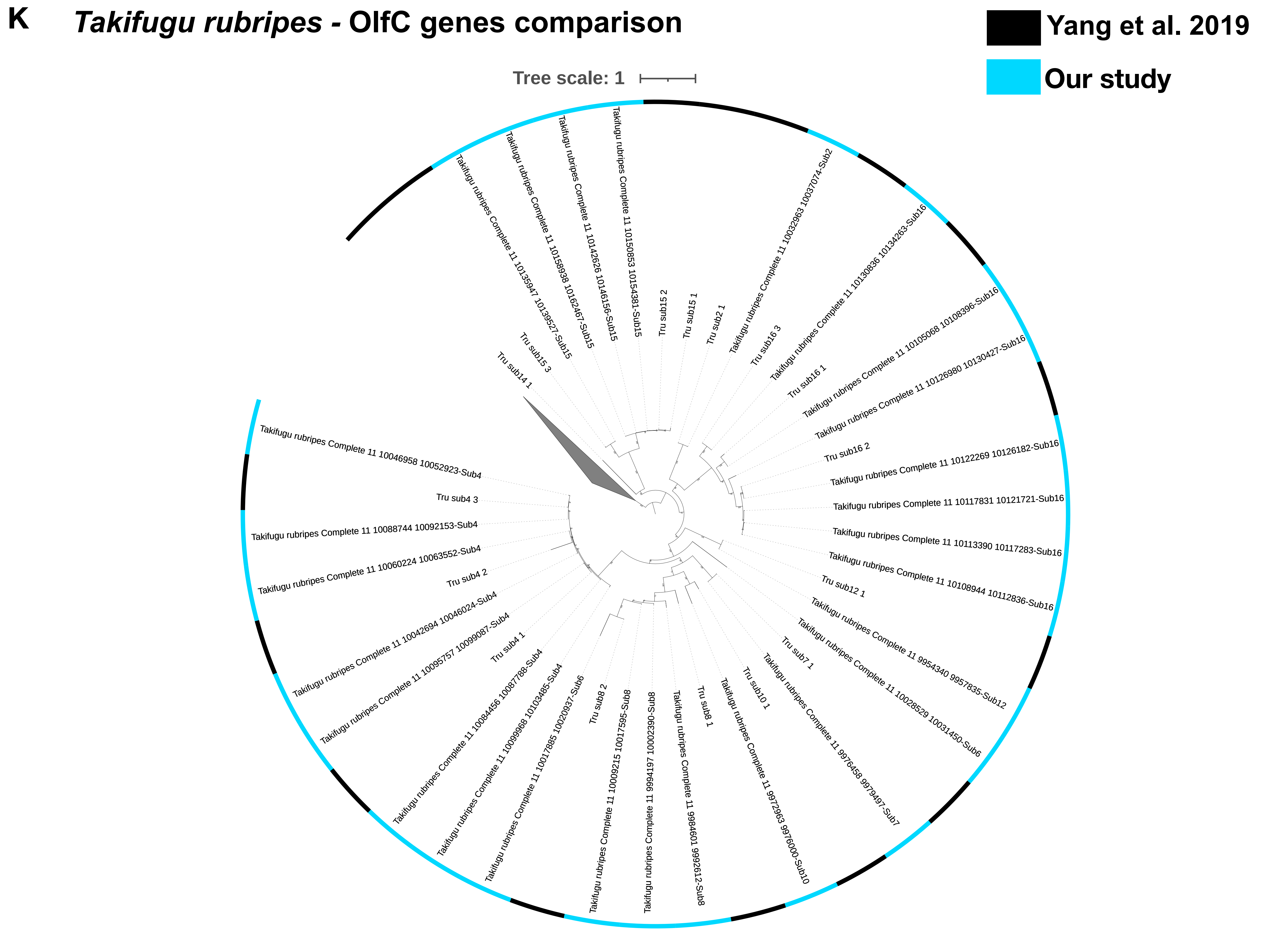
K
Takifugu rubripes - OlfC genes comparison
Yang et al. 2019
Our study
Tree scale: 1
Tru sub14 1
Tru sub15 3
Takifugu rubripes Complete 11 10135947 10139527-Sub15
Takifugu rubripes Complete 11 10158938 10162467-Sub15
Takifugu rubripes Complete 11 10142626 10146156-Sub15
Takifugu rubripes Complete 11 10150853 10154381-Sub15
Tru sub15 2
Tru sub15 1
Tru sub2 1
Takifugu rubripes Complete 11 10032963 10037074-Sub2
Tru sub16 3
Takifugu rubripes Complete 11 10130836 10134263-Sub16
Tru sub16 1
Takifugu rubripes Complete 11 10105068 10108396-Sub16
Takifugu rubripes Complete 11 10126980 10130427-Sub16
Tru sub16 2
Takifugu rubripes Complete 11 10122269 10126182-Sub16
Takifugu rubripes Complete 11 10117831 10121721-Sub16
Takifugu rubripes Complete 11 10113390 10117283-Sub16
Takifugu rubripes Complete 11 10108944 10112836-Sub16
Tru sub12 1
Takifugu rubripes Complete 11 9954340 9957835-Sub12
Takifugu rubripes Complete 11 10028529 10031450-Sub6
Tru sub7 1
Takifugu rubripes Complete 11 9976458 9979497-Sub7
Tru sub10 1
Takifugu rubripes Complete 11 9972963 9976000-Sub10
Tru sub8 1
Takifugu rubripes Complete 11 9984601 9992612-Sub8
Takifugu rubripes Complete 11 9994197 10002390-Sub8
Takifugu rubripes Complete 11 10009215 10017595-Sub8
Tru sub8 2
Takifugu rubripes Complete 11 10017885 10020937-Sub6
Takifugu rubripes Complete 11 10099968 10103485-Sub4
Takifugu rubripes Complete 11 10084456 10087788-Sub4
Tru sub4 1
Takifugu rubripes Complete 11 10095757 10099087-Sub4
Takifugu rubripes Complete 11 10042694 10046024-Sub4
Tru sub4 2
Takifugu rubripes Complete 11 10060224 10063552-Sub4
Takifugu rubripes Complete 11 10088744 10092153-Sub4
Tru sub4 3
Takifugu rubripes Complete 11 10046958 10052923-Sub4

L

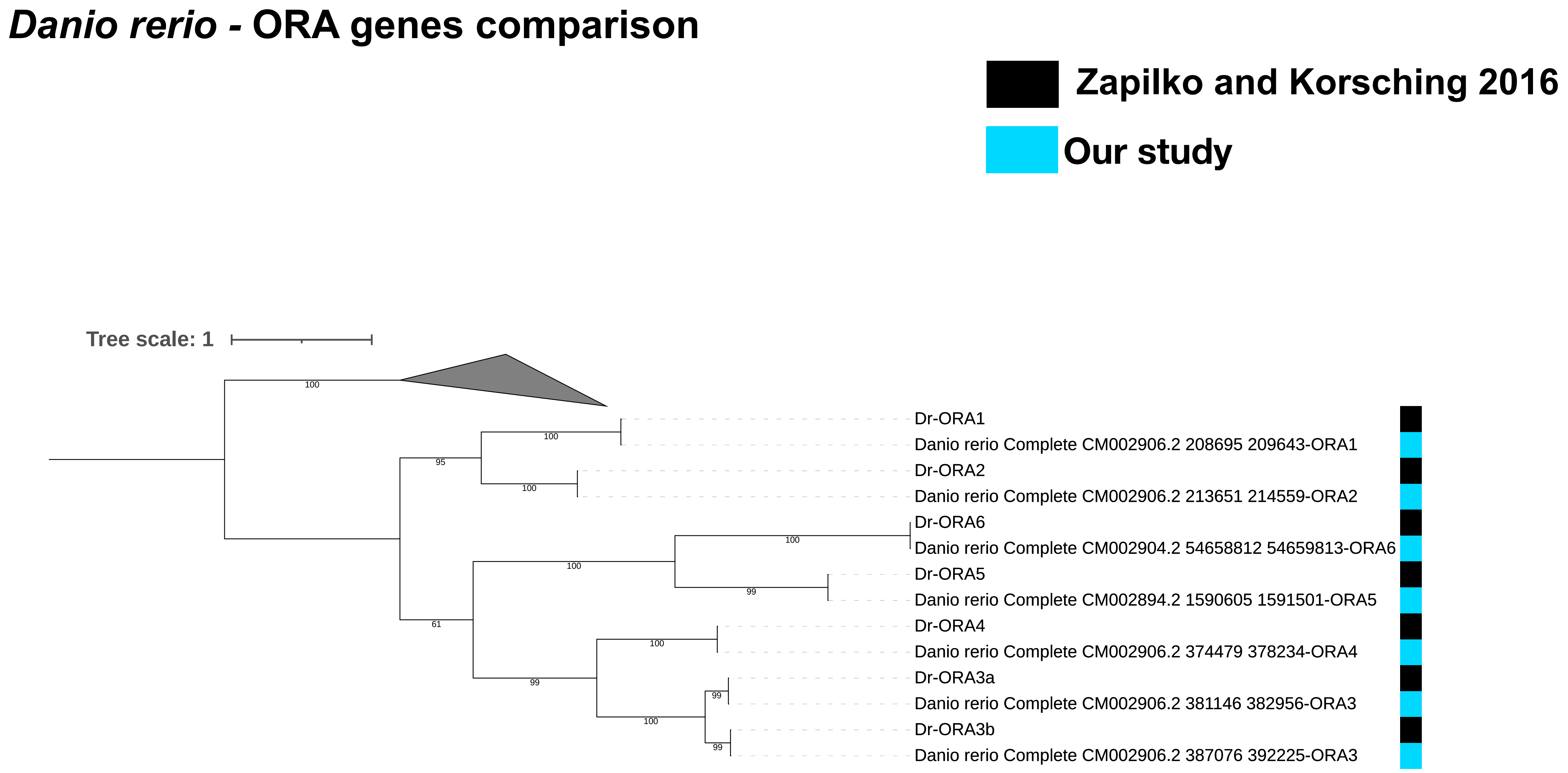
Danio rerio - ORA genes comparison
Zapilko and Korsching 2016
Our study
Tree scale: 1
100
100
95
100
100
100
99
61
100
99
99
100
99
Dr-ORA1
Danio rerio Complete CM002906.2 208695 209643-ORA1
Dr-ORA2
Danio rerio Complete CM002906.2 213651 214559-ORA2
Dr-ORA6
Danio rerio Complete CM002904.2 54658812 54659813-ORA6
Dr-ORA5
Danio rerio Complete CM002894.2 1590605 1591501-ORA5
Dr-ORA4
Danio rerio Complete CM002906.2 374479 378234-ORA4
Dr-ORA3a
Danio rerio Complete CM002906.2 381146 382956-ORA3
Dr-ORA3b
Danio rerio Complete CM002906.2 387076 392225-ORA3

**M** ***Gasterosteus aculeatus* - ORA genes comparison**

Zapilko and Korsching 2016

Our study

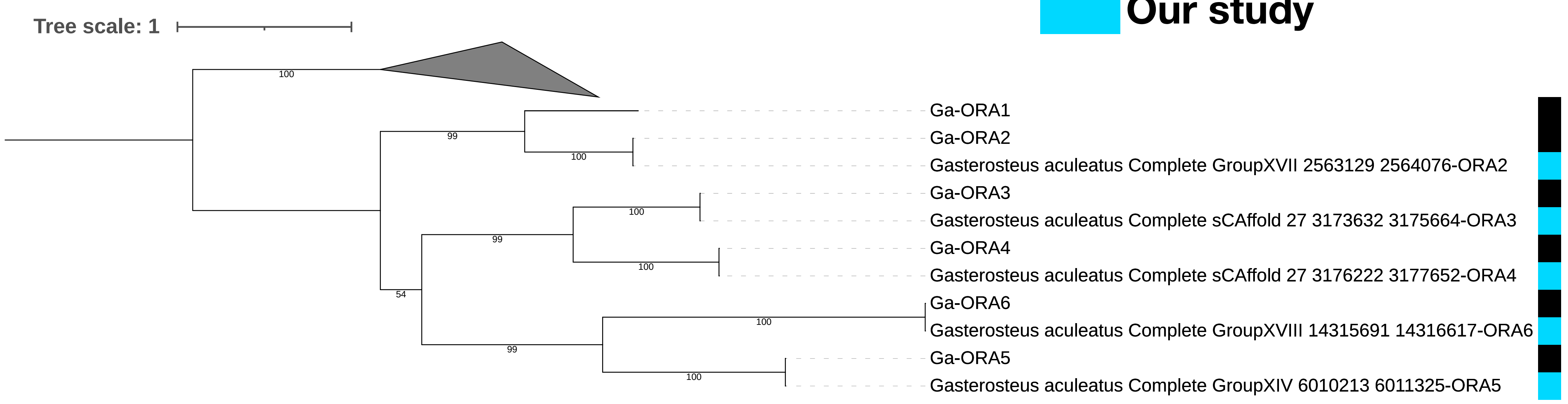


ORA1 is found as pseudogene (insertion of 2bp) in the current stickleback genome assembly (BROADS1)

```
C4 Alignment:
------------
         Query: Ga-ORA1
        Target: GroupXVII:2554871-2563367 [revcomp]
         Model: protein2genome:bestfit
     Raw score: 1276
   Query range: 0 -> 313
  Target range: 4958 -> 4001

    1 : MetAspLeuCysValThrIleLysGlyValSerPheLeuLeuGlnThrGlyMetGlyIleLe :   21
        ||||||||||||||||||||||||||||||||||||||||||||||||||||||||||||||
        MetAspLeuCysValThrIleLysGlyValSerPheLeuLeuGlnThrGlyMetGlyIleLe
 4958 : ATGGATCTGTGCGTCACCATCAAAGGGGTCTCCTTCCTCCTGCAAACAGGCATGGGCATCTT : 4898

   22 : uGlyAsnThrValValLeuLeuAlaTyrAlaGlnLeuIleTyrAlaGluProLysLeuLeuP :   42
        ||||||||||||||||||||||||||||||||||||||||||||||||||||||||||||||
        uGlyAsnThrValValLeuLeuAlaTyrAlaGlnLeuIleTyrAlaGluProLysLeuLeuP
 4897 : AGGGAACACGGTGGTGCTGCTGGCCTACGCTCAGCTCATCTACGCCGAGCCCAAGCTCCTAC : 4835

   43 : roValAspMetIleLeuCysHisLeuAlaPheAlaAsnLeuMetLeuLeuLeuThrArgCys :   62
        ||||||||||||||||||||||||||||||||||||||||||||||||||||||||||||||
        roValAspMetIleLeuCysHisLeuAlaPheAlaAsnLeuMetLeuLeuLeuThrArgCys
 4834 : CCGTGGACATGATCCTGTGCCACCTGGCCTTCGCCAACCTGATGCTGCTGCTGACCCGCTGC : 4775

   63 : ValProGlnThrMetSerValPheGlyLeuArgAspLeuLeuGlyAspProGlyCysLysVa :   83
        ||||||||||||||||||||||||||||||||||||||||||||||||||||||||||||||
        ValProGlnThrMetSerValPheGlyLeuArgAspLeuLeuGlyAspProGlyCysLysVa
 4774 : GTCCCGCAGACCATGAGCGTGTTCGGGCTGAGGGACCTGCTGGGTGACCCCGGCTGCAAGGT : 4712

   84 : lValIleTyrAlaTyrArgIleGlyArgAlaLeuSerValCysValThrCysMetLeuSerV :  104
        ||||||||||||||||||||||||||||||||||||||||||||||||||||||||||||||
        lValIleTyrAlaTyrArgIleGlyArgAlaLeuSerValCysValThrCysMetLeuSerV
 4711 : GGTGATCTACGCCTACCGCATCGGCCGGGCTTTGTCGGTCTGCGTCACCTGCATGCTCAGCG : 4649

  105 : alPheGlnAlaValThrLeuAlaPro--AlaGlyProArgLeuSerArgLeuLysProAlaL :  124
        ||||||||||||||||||||   |||##||||||||||||||||||||||||||||||||||
        alPheGlnAlaValThrLeu---Pro##AlaGlyProArgLeuSerArgLeuLysProAlaL
 4648 : TCTTTCAGGCGGTGACCTTG---CCCTCGCCGGACCCCGTCTGTCACGGTTGAAGCCCGCAC : 4590
```

N

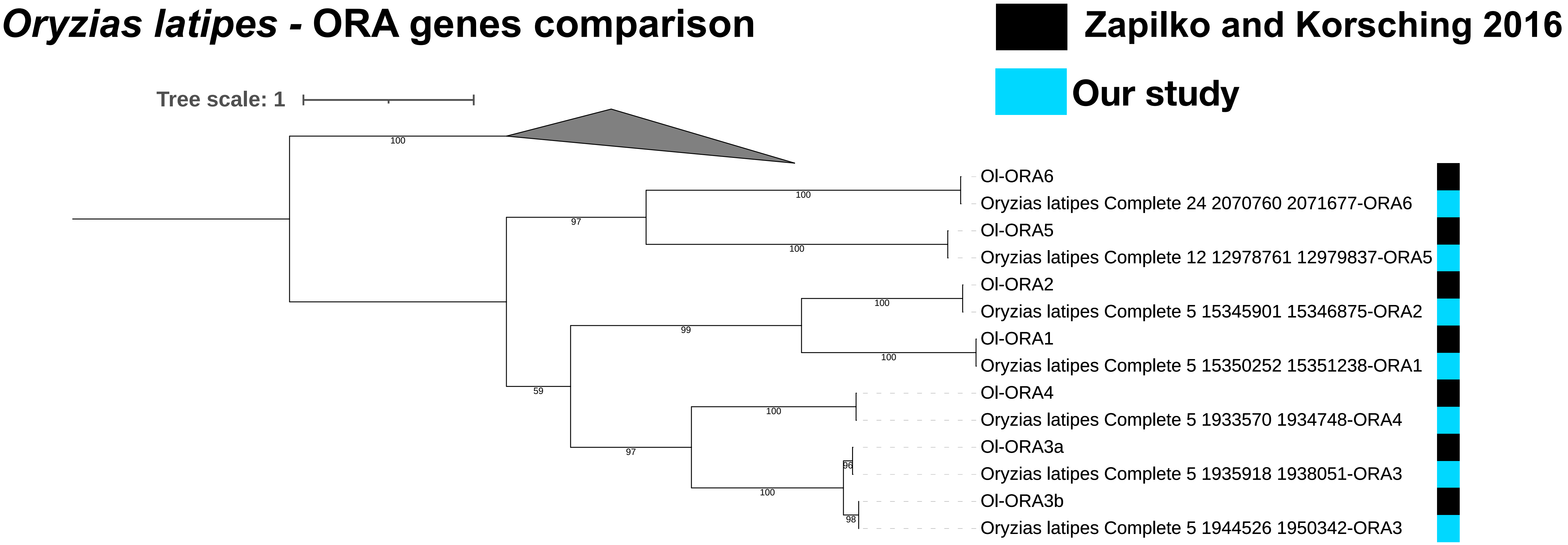
Oryzias latipes - ORA genes comparison
Zapilko and Korsching 2016
Our study
Tree scale: 1
Ol-ORA6
Oryzias latipes Complete 24 2070760 2071677-ORA6
Ol-ORA5
Oryzias latipes Complete 12 12978761 12979837-ORA5
Ol-ORA2
Oryzias latipes Complete 5 15345901 15346875-ORA2
Ol-ORA1
Oryzias latipes Complete 5 15350252 15351238-ORA1
Ol-ORA4
Oryzias latipes Complete 5 1933570 1934748-ORA4
Ol-ORA3a
Oryzias latipes Complete 5 1935918 1938051-ORA3
Ol-ORA3b
Oryzias latipes Complete 5 1944526 1950342-ORA3

O

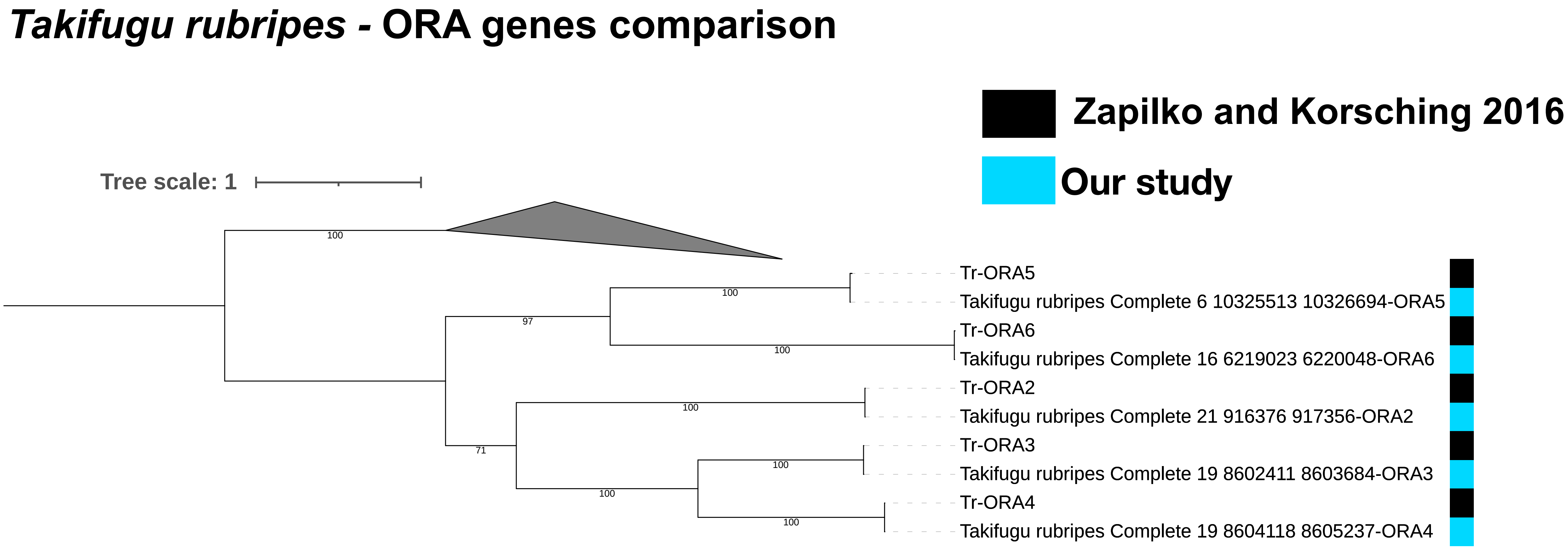
Takifugu rubripes - ORA genes comparison
Zapilko and Korsching 2016
Our study
Tree scale: 1
Tr-ORA5
Takifugu rubripes Complete 6 10325513 10326694-ORA5
Tr-ORA6
Takifugu rubripes Complete 16 6219023 6220048-ORA6
Tr-ORA2
Takifugu rubripes Complete 21 916376 917356-ORA2
Tr-ORA3
Takifugu rubripes Complete 19 8602411 8603684-ORA3
Tr-ORA4
Takifugu rubripes Complete 19 8604118 8605237-ORA4

**P** ***Polypterus senegalus*** **- OR genes comparison**

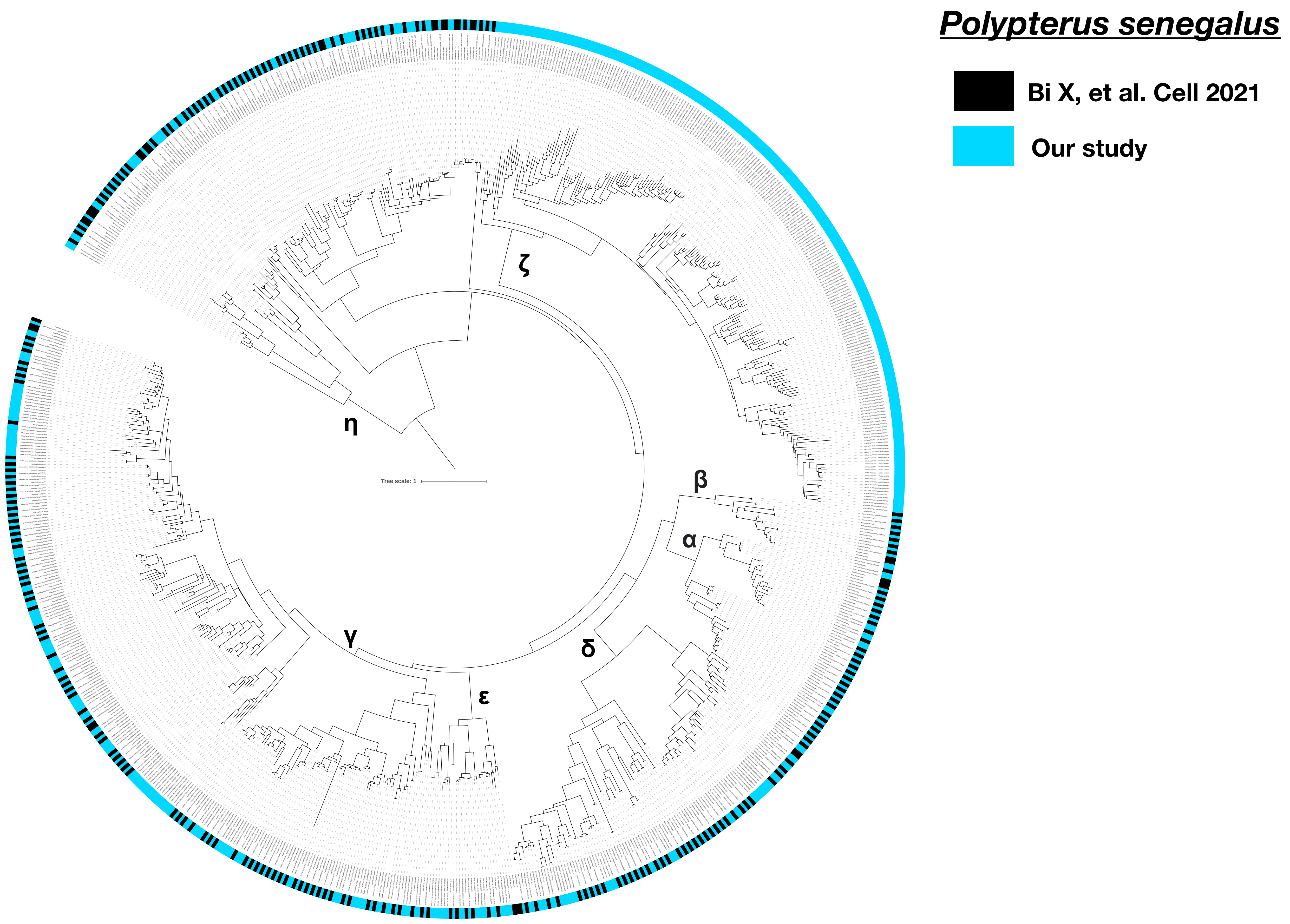
