## Supplementary figures and images for "Coevolution of the olfactory organ and its receptor repertoire in ray-finned fishes"

### Supplementary data 3

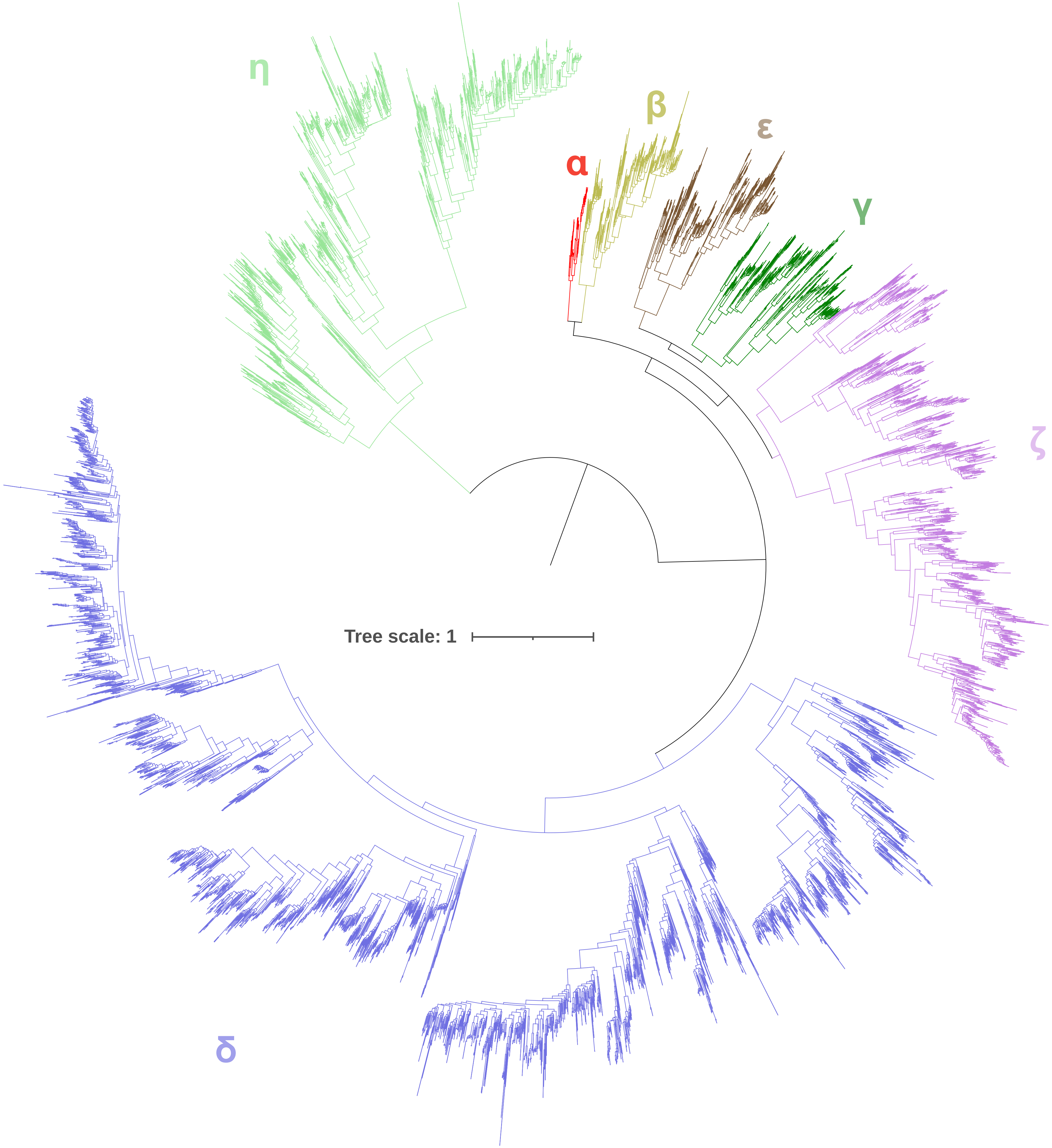

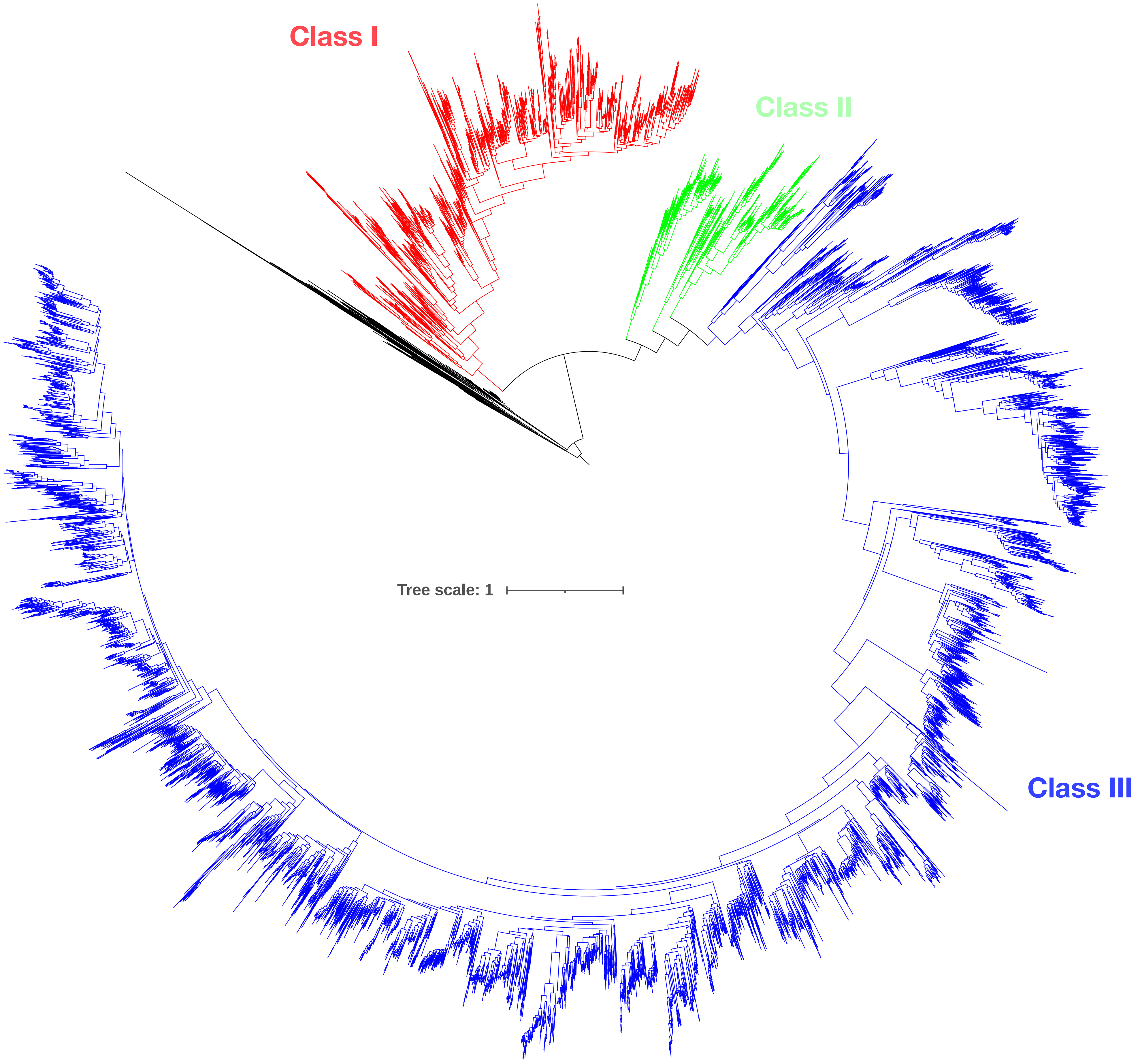

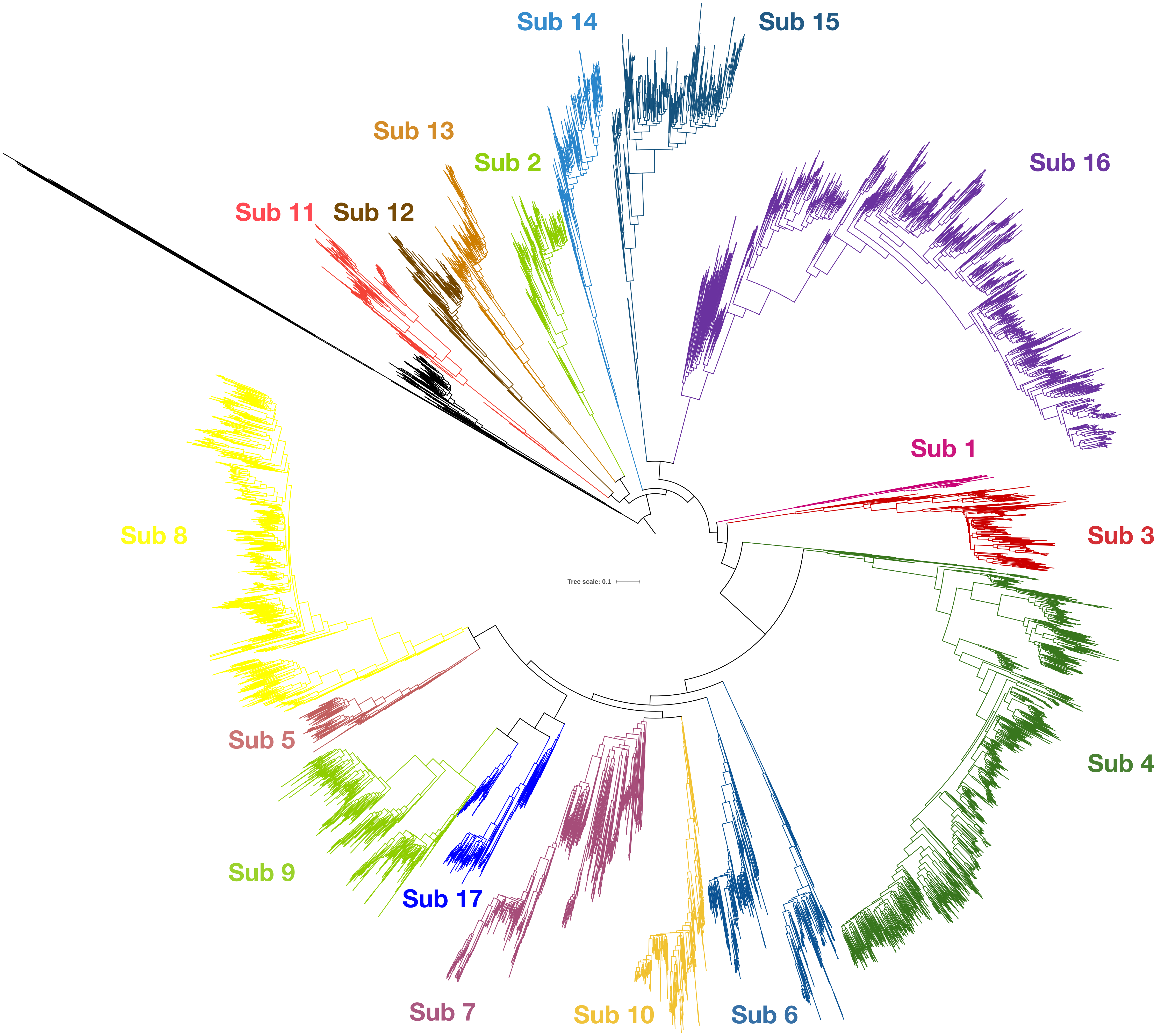

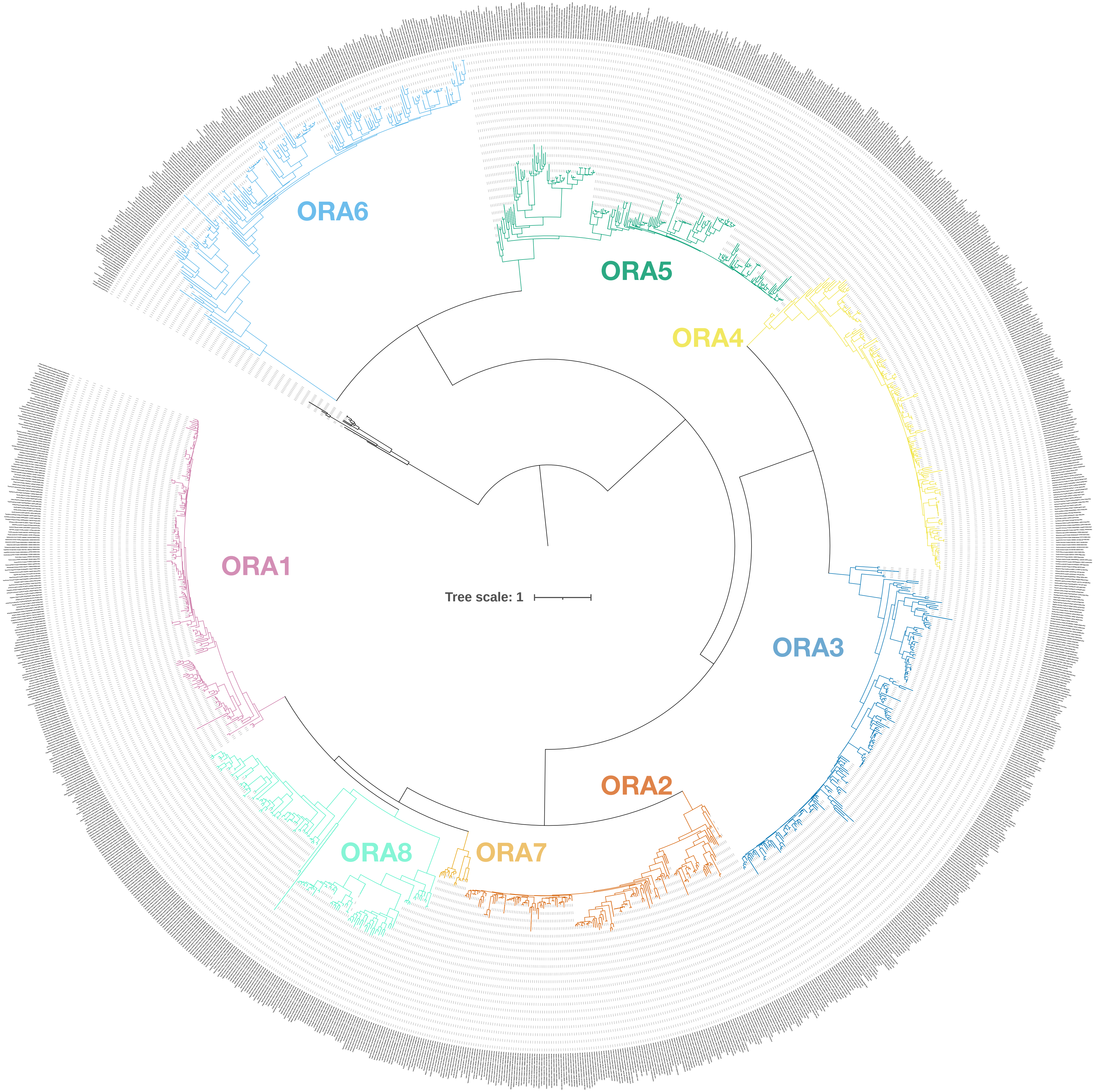
